# Meristem dormancy in a dichotomous branching system is regulated by a liverwort-specific miRNA and a clade III *SPL* gene in *Marchantia polymorpha*

**DOI:** 10.1101/2022.10.03.510622

**Authors:** Susanna Streubel, Sebastian Deiber, Johannes Rötzer, Magdalena Mosiolek, Katharina Jandrasits, Liam Dolan

## Abstract

The shape of modular organisms depends on branching architecture, which in plants is determined by the fates of generative centres called meristems. The branches of the liverwort *Marchantia polymorpha* are derived from two adjacent meristems that develop at thallus apices. These meristems may be active and develop branches or may be dormant and do not form branches. The relative number and position of active and dormant meristems defines overall shape and form of the thallus. We show that the clade III SQUAMOSA PROMOTER BINDING PROTEIN LIKE (SPL) transcription factor, Mp*SPL1*, is required for meristem dormancy. The activity of Mp*SPL1* is regulated by the liverwort-specific Mpo-MR13 miRNA which in turn is regulated by *PIF*-mediated phytochrome signaling. An unrelated miRNA, MIR156, represses a different *SPL* gene (belonging to clade IV) that inhibits branching during the shade avoidance response in *Arabidopsis thaliana*. This suggests that a conserved mechanism of phytochrome signaling modulates branching architecture in liverworts and angiosperms and therefore likely operated in the last common ancestor. However, *PIF*-mediated phytochrome signaling represses the expression of different miRNA genes with different *SPL* targets during dichotomous, apical branching in liverworts and during lateral, subapical branching in angiosperms. We speculate that the mechanism that acts downstream of light and regulates meristem dormancy evolved independently in liverworts and angiosperms.

## INTRODUCTION

Modular organisms – plants, colonial animals, and fungi – grow through the proliferation of fundamental units of organization called modules (White, 1979; Bell et al., 1986; Harper et al., 1986). Modules are connected to each other in branches and the ultimate shape of the organism depends on the three-dimensional organization of the branching network that connects individual modules. In extant land plants, branching axes comprise the shoot and root system of the diploid stage of vascular plants while the small haploid phase of the life cycle does not branch (Bierhorst, 1971). Branching axes occur in the haploid stage of the life cycle of the bryophytes, the other monophyletic group that makes up the land plants, but there is no branching in the diploid phase. Phylogenetics and comparative genomics suggest that branching axes occurred in both the haploid and diploid phases of the life cycle in the last common ancestor of the extant land plants (Nishiyama et al., 2004; Puttick et al., 2018; Harris et al., 2020; Clark et al., 2022). Subsequently, branching was lost in the diploid phase of the bryophytes and in the haploid phase of the vascular plants as exemplified by the seed plants (Puttick et al., 2018; Harris et al., 2020).

There are two classes of branching in land plants (Kenrick & Crane, 1997; Hetherington et al., 2020). Dichotomous branching is a form of apical branching that occurs when an apical meristem duplicates and each daughter meristem has the potential to form a branch (Bierhorst, 1971; Gola et al., 2014). It occurs in bryophyte gametophytes and lycophyte sporophytes (Schuster, 1984a; Gola et al., 2014). Lateral branching is a form of subapical branching, where new meristems are formed in the axils of determinate structures (leaves) and can form branches. Lateral branching is found in euphyllophyte sporophytes. The haploid body of *Marchantia polymorpha* consists of a bifurcating system of thallus branches which develop from pairs of meristems located at thallus apices (Shimamura, 2016; Solly et al., 2017).

Here, we demonstrate that the architecture of the dichotomously branching *M. polymorpha* thallus is determined by the relative number of active and dormant meristems formed at thallus apices. Meristem dormancy is controlled by the activity of the clade III SQUAMOSA PROMOTER BINDING PROTEIN LIKE gene Mp*SPL1*, a transcription factor that is required for meristem dormancy. The activity of Mp*SPL1* is inhibited by the liverwort-specific Mpo-MR13 miRNA whose expression is regulated by phytochrome signaling to modulate meristem dormancy and in turn branching architecture.

## RESULTS

### Meristem dormancy modulated by the ratio of red and far-red light determines the branching architecture of the *Marchantia polymorpha* thallus network

The haploid phase of the *M. polymorpha* life cycle comprises a radially symmetrical network of bifurcating branches (Solly et al., 2017). New branches develop from meristems that are formed by the synchronised duplication of apical meristems. Each apical meristem duplication event doubles the number of meristems and has the potential to double the number of branches. If each daughter meristem developed as a branch with each order of branching, the resulting exponential increase in the number of branches would lead to self-shading. We hypothesized that a mechanism to suppress self-shading exists by inducing dormancy (repressing the development) of meristems formed upon apical meristem bifurcation. Furthermore, angiosperms avoid self-shading by imposing meristem dormancy which is promoted by a low ratio of red to far-red (R:FR) ambient light. Therefore, we predicted that the *M. polymorpha* branching network avoids self-shading by inducing dormancy in some meristems and that dormancy is regulated by the R:FR ratio of ambient light.

To test if the R:FR ratio in ambient light modulates meristem dormancy in developing *M. polymorpha* thalli, we grew Tak-1 and Tak-2 accessions in two different R:FR light regimes – R:FR of 1.65 and R:FR of 0.29 – for 15 days (after an initial exposure to high R:FR light (R:FR = 1.65) for 12 days) (Figure 1A). In plants grown in high R:FR light, each meristem that formed by apical meristem duplication during the first four plastochrons remained active and formed an axis (Figure 1B-D). There was no meristem dormancy (Figure 1D) and therefore no evidence of apical dominance during the first four plastochrons in these high R:FR (1.65) conditions. This growth pattern resulted in the formation of a radially bifurcating thallus network.

**Figure 1:**
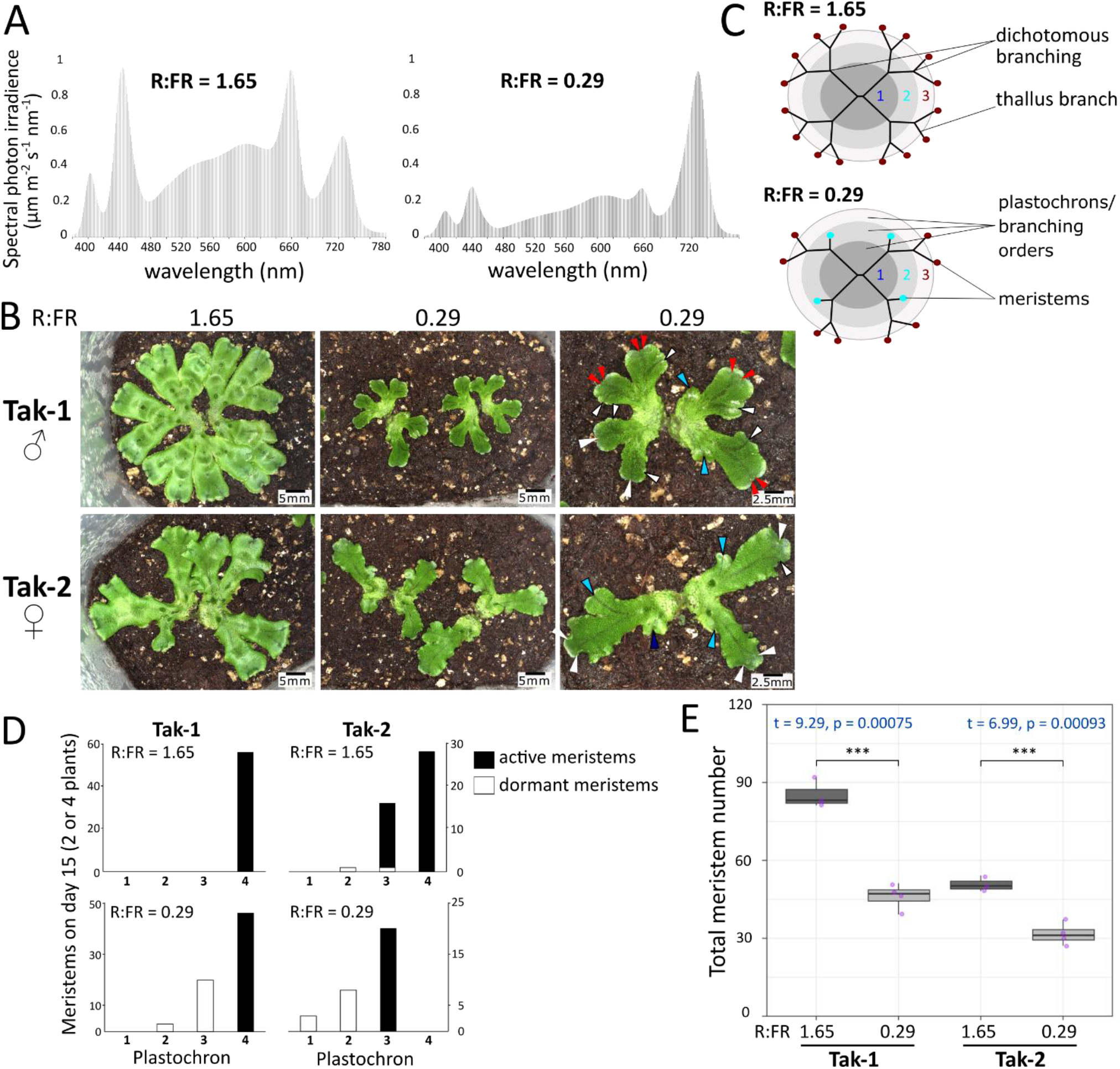
*M. polymorpha* thallus develops apical dominance in low R:FR light. **(A)** Spectral photon irradiance in growth chambers with high or low R:FR ratio as measure with the MK350S Premium Spectrometer. R:FR ratios were calculated using the formula 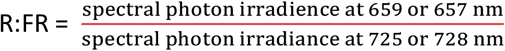 **(B)** Wild-type gemmae were grown in constant white light (R:FR = 1.65) for 12 days, then either kept under white light or transferred to far-red enriched illumination (R:FR = 0.29) for 15 more days. Plants in the two right-hand panels are higher magnifications of plants from the middle panels with arrow heads indicating branching oders (bo): blue – bo 1, light blue – bo 2, white – bo 3, red – bo 4. **(C)** Schematics illustrating thallus branching patterns in high (R:FR = 1.65) and low (R:FR = 0.29) R:FR light, with black lines indicating thallus branches, coloured dots representing thallus meristems and large grey circles indicating plastochrons. Meristems are in plastochrons two (turquoise dots) and three (maroon dots). **(D)** Absolute number of active meristems (black bars) and dormant meristems (white bars) in different plastochrons on day 15 of growth in high or low R:FR light as described in (B). For each line, Tak-1 and Tak-2, we counted and evaluated all meristems from two plants grown in high R:FR light and from four plants grown in low R:FR light. **(E)** Average total number of meristems per plant grown in high and low R:FR light. Significant differences were calculated using Welch’s t-test with n = 3 and n = 4, respectively.

In plants grown in low R:FR light (R:FR = 0.29) for 15 days, one of the two meristems formed by meristem duplication became dormant soon after initiation while the other was active and developed an axis (Figure 1B-D). Repressed, dormant branches were arranged on alternate sides of the axis at each branching event; if, at plastochron n, the daughter meristem on the left side of the thallus axis remained active and the right daughter meristem was repressed, then at plastochron n+1 the daughter meristem on the right side of the axis was active and the daughter meristem on the left was repressed (Figure 6A). The development of dormant meristems after each meristem duplication resulted in the formation of linear axes without branching (a branching event requires both daughter meristems to develop axes) (Figure 1B). Because of the impact of meristem dormancy on network architecture and the reduction in the number of axes with active meristems, the total number of meristems that formed in low F:FR (0.29) was approximately half the number that formed in high R:FR (1.65) (Figure 1E). We conclude that one of the two daughter meristems produced by apical meristem duplication often becomes dormant in low R:FR light while both meristems remain active and form a branch in high R:FR light. The demonstration that ambient light controls branching by regulating meristem dormancy, as it does in angiosperm apical dominance, leads to the hypothesis that apical dominance modulates branching of the *M. polymorpha* thallus network.

### Mp*PHY* and Mp*PIF* antagonistically regulate meristem dormancy and thallus branching

Since meristem dormancy in *M. polymorpha* is modulated by the R:FR ratio of ambient light, we hypothesized that phytochrome signalling would modulate meristem dormancy during the development of the thallus. Red and far-red light are perceived by phytochrome (PHY) photoreceptors that modulate the activity of PHYTOCHROME-INTERACTING FACTOR (PIF) transcriptional regulators. In *M. polymorpha*, the relative activities of MpPHY and MpPIF are determined by the R:FR light ratio (Inoue et al., 2016). To test if Mp*PHY* regulates far-red light-modulated meristem dormancy, we generated two loss of function (lof) mutant alleles in Mp*PHY* using CRISPR/Cas9 (Suppl. Figure 1). There was a constitutive far-red light response in Mp*phy-27*^*lof*^ and Mp*phy-38*^*lof*^ mutants that lacked the chromophore binding domain of wild-type MpPHY. Mp*phy-27*^*lof*^ and Mp*phy-38*^*lof*^ mutant plants were pale green while wild-type is dark green (Figure 2A). Mp*phy-27*^*lof*^ and Mp*phy-38*^*lof*^ mutant thalli bent upward when grown in either high or low R:FR, a phenotype that only develops in wild-type plants when grown in low R:FR (Suppl. Figure2). These phenotypes demonstrated that the R:FR light response was defective in the Mp*phy-27*^*lof*^ and Mp*phy-38*^*lof*^ mutants. We then tested the hypothesis that apical dominance would be stronger in Mp*phy-27*^*lof*^ and Mp*phy-38*^*lof*^ than in wild-type plants in both high R:FR and low R:FR light because of the constitutive far-red light response resulting from the lack of functional Mp*PHY* in these mutants. Mp*phy-27*^*lof*^, Mp*phy-38*^*lof*^ and wild-type were grown in either high R:FR or low R:FR light for 21 days following initial growth in high R:FR light for 14 days, and the branching phenotypes were analysed. The two Mp*phy*^*lof*^ mutants developed high levels of meristem dormancy in both high and low R:FR light while wild-type developed meristem dormancy in low R:FR but not in high R:FR (Figure 2A, 2C, 2F). Both Mp*phy*^*lof*^ mutants developed linear, dominant thallus axes without forming a branching network. These axes were linear because from the first plastochron, one daughter meristem became dormant soon after meristem duplication while the other developed the dominant thallus axis (Figure 2C, 2F). The frequency of meristem dormancy was higher in Mp*phy*^*lof*^ mutants than in wild-type in both high R:FR and low R:FR light. Consequently, total meristem number, which included active and dormant meristems, was less in the Mp*phy*^*lof*^ mutants than in wild-type (Figure 2B). These data are consistent with the hypothesis that active Mp*PHY* is required for branching and the repression of meristem dormancy in high R:FR light.

**Figure 2:**
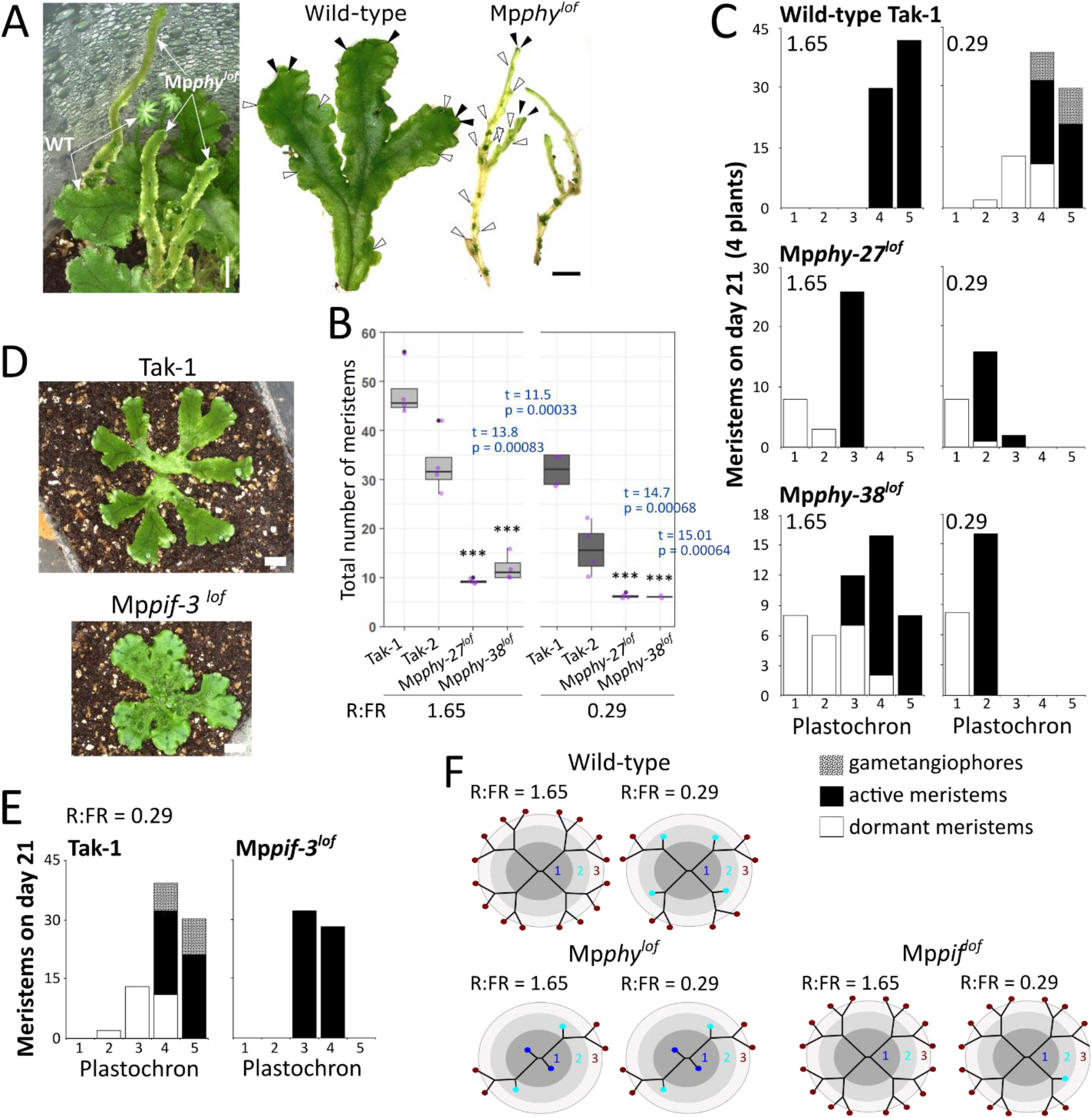
Mp*PHY* and Mp*PIF* control meristem dormancy and apical dominance in an antagonal manner. **(A)** 4-months-old Mp*phy*^*lof*^ mutant next to Tak-2 wild-type grown in low R:FR (0.29) in a pot (left image), and branches removed from these plants (right image). The wild-type plant is at least two months younger than the Mp*phy*^*lof*^ plants. Black and white arrow heads indicate active and dormant meristems. Scale bars are 5mm. **(B)** Average total number of meristems per plant in Tak-1 and Tak-2 wild-types and Mp*phy*^*lof*^ mutants grown in high and low R:FR light. Significant differences between the male mutants and Tak-1 were calculated using Welch’s t-test with n = 4 (n = 2 for Mp*phy-38*^*lof*^ in R:FR = 0.29). **(C)** Absolute number of active meristems (black bars) and dormant meristems (white bars) in different plastochrons in Tak-1 and male Mp*phy*^*lof*^ mutants. Plants were grown in high or low R:FR for 21 days after 14 days of pre-growth in high R:FR light. 4 plants were evaluated per mutant line and light condition, 3 plants for Tak-1 per light condition. By day 21, Tak-1 developed gametangiophores in low R:FR light (patterned fractions of the bars). **(D)** Mp*pif-3*^*lof*^ mutant and Tak-1 wild-type gemmae grown in high R:FR for 14 days, then in low R:FR for 21 days. Scale bars are 5mm. Mp*pif-3*^*lof*^ thallus grows flat on the surface of the soil, while Tak-1 thallus bends hyponastically towards the light source. **(E)** Absolute number of active meristems (black bars) and dormant meristems (white bars) in different plastochrons in Tak-1 (3 plants) and Mp*pif-3*^*lof*^ mutant (5 plants) grown in low R:FR as described in (D). **(F)** Schematics illustrating thallus morphologies of wild-type, Mp*phy*^*lof*^ and Mp*pif*^*lof*^ mutants in high (1.65) and low (0.29) R:FR. Labelling of the schematics is the same as in Figure 1.

Since Mp*phy*^*lof*^ mutants developed more meristem dormancy than wild-type and MpPHY represses the function of MpPIF (Inoue et al., 2016), we hypothesized that Mp*pif*^*lof*^ mutants would develop less meristem dormancy than wild-type. To determine the function of Mp*PIF* in branching and meristem dormancy, we generated a Mp*pif*^*lof*^ mutant using CRISPR/Cas9 (Suppl. Figure 2). To verify that Mp*PIF* function was defective in Mp*pif-3*^*lof*^ mutants, we assessed the orientation of mutant thalli grown in low R:FR. Consistent with defective Mp*PIF* function, Mp*pif-3*^*lof*^ thalli grew horizontally in low R:FR light while wild-type thalli bent upward (Figure 2D, Suppl. Figure 3). To determine if Mp*PIF*-mediated signalling is required for meristem dormancy in low R:FR, we grew the Mp*pif-3*^*lof*^ mutant in low R:FR light (after initial growth in high R:FR for 14 days) and characterised its branching phenotype after 21 days (Figure 2D-F). While meristem dormancy developed in wild-type plants in low R:FR light there was no meristem dormancy in Mp*pif-3*^*lof*^ mutants (Figure 2E, F). All meristems that formed by apical meristem duplication developed into thallus branches in the mutant in low R:FR light, which meant that all meristems in mutant plants were in plastochron 3 or 4 and active at the time of observation while in wild-type, some meristems that developed in plastochrons 2, 3 and 4 were dormant (Figure 2E). The apical dominance phenotype of the Mp*pif-3*^*lof*^ mutant grown in low R:FR light was indistinguishable from wild-type grown in high R:FR light (Figure 2F). We conclude that Mp*PIF* promotes meristem dormancy in plants grown in low R:FR light.

**Figure 3:**
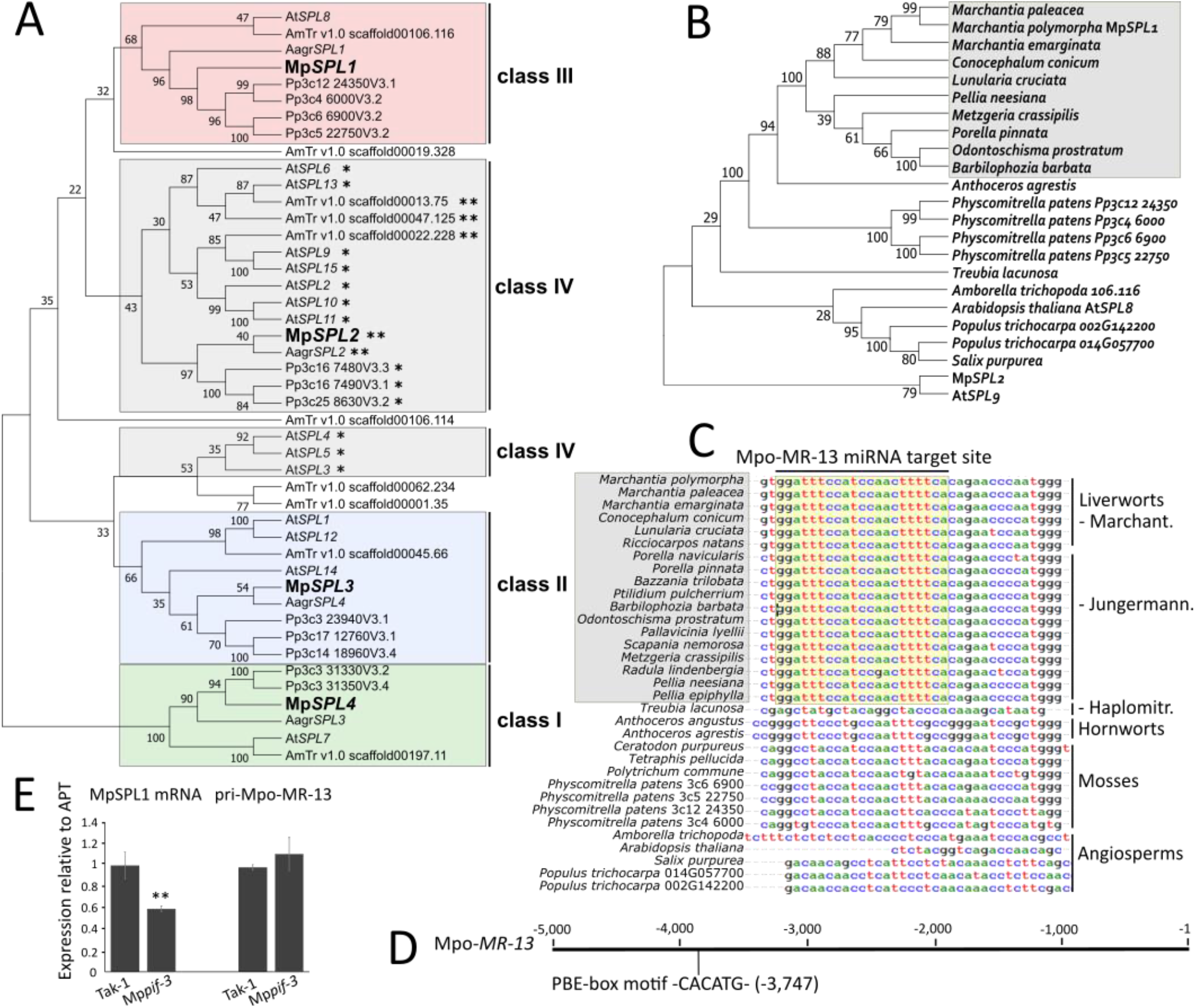
Class III Mp*SPL1* is target of the liverwort-specific miRNA Mpo-MR-13, which is regulated by phytochrome. **(A)** Unrooted cladogram of *SPL* genes from angiosperms *Arabidopsis thaliana* (At) and *Amborella trichopoda* (Amtr), the moss *Physcomitrella patens* (Pp), the hornwort *Anthoceros agrestis* (Aagr) and the liverwort *Marchantia polymorpha* (Mp). Branch support values are bootstrap values from a maximum likelihood analysis with 500 resampling steps conducted in Mega-X (Kumar et al., 2018). The underlying MSA generated by the Mafft online programme was manually trimmed down to the SBP box domain and conserved C-terminal sequences. *SPL* genes marked by asterisks are known targets of miR156* or miR529c** miRNAs. **(B)** Clade III *SPL* gene tree with species from (C) including a subset of liverwort species, generated from a trimmed MSA of amino acid sequences. Maximum likelihood analysis included 1000 bootstrap resampling steps. At*SPL9* and Mp*SPL2* form an outgroup, branch support values are bootstrap values. **(C)** Segment of a MSA of clade III *SPL* gene sequences from liverworts, hornworts, mosses and angiosperms around the Mpo-MR-13 miRNA binding site highlighted in yellow. **(D)** Putative MpPIF binding site (5′-CACATG-3′) in the genomic region 5kb upstream of the Mpo-MR-13 miRNA sequence (TGAAAAGTGGGATGGAGATCC). **(E)** Steady state levels of MpSPL1 mRNA and pri-Mpo-MR-13 pri-miRNA in Tak-1 and Mp*pif-3*^*lof*^ mutant gemmalings grown in low R:FR light for 13 days. Significant differences were calculated using Student’s t-test with p<0.05*, p<0.01** and p<0.001***

Taken together, these data indicate that Mp*PHY* inhibits meristem dormancy and promotes branching in high R:FR, while Mp*PIF* promotes meristem dormancy and represses branching in low R:FR. These data are consistent with the hypothesis that meristem dormancy is modulated by the ratio of R:FR light and regulated by phytochrome signalling.

### Mpo-MR-13 is a liverwort specific miRNA that targets the mRNA of clade III Mp*SPL1*

We set out to identify the molecular mechanism by which phytochrome signalling controls meristem dormancy during thallus development. While our data demonstrate that phytochrome signalling modulates meristem dormancy, low R:FR light also induces the development of gametangiophores – the reproductive structures that produce egg or sperm – in *M. polymorpha* by promoting Mp*SPL2* expression. miRNA529c represses Mp*SPL2* function and determines the time at which gametangiophores develop in *M. polymorpha* (Tsuzuki et al., 2019). We hypothesized that phytochrome signalling mediated by Mp*PIF* would modulate miRNA529c expression. Consistent with this hypothesis, we found a PIF-binding motif (PBE-box motif) (Leivar & Monte, 2014) 2,572 bp upstream of the genomic miR529c sequence suggesting that Mp*PIF* controls the expression of this gene (Suppl. Figure 4). Furthermore, steady state levels of MpSPL2 mRNA, the target of miR529c, were lower in the Mp*pif-3*^*lof*^ mutant than in wild type after 14 days of growth in low R:FR light where Mp*PIF*-mediated repression of miR529c is active in wild-type (Suppl. Figure 5). This suggests that in low R:FR light, Mp*PIF* inhibits the expression of miR529c leading to an increase in steady state MpSPL2 mRNA levels. Since MpSPL2 promotes the development of gametangiophores but not meristem dormancy (Tsuzuki et al., 2019), we set out to test the hypothesis that Mp*PIF* regulates a different miRNA-Mp*SPL* gene pair to modulate meristem dormancy.

**Figure 4:**
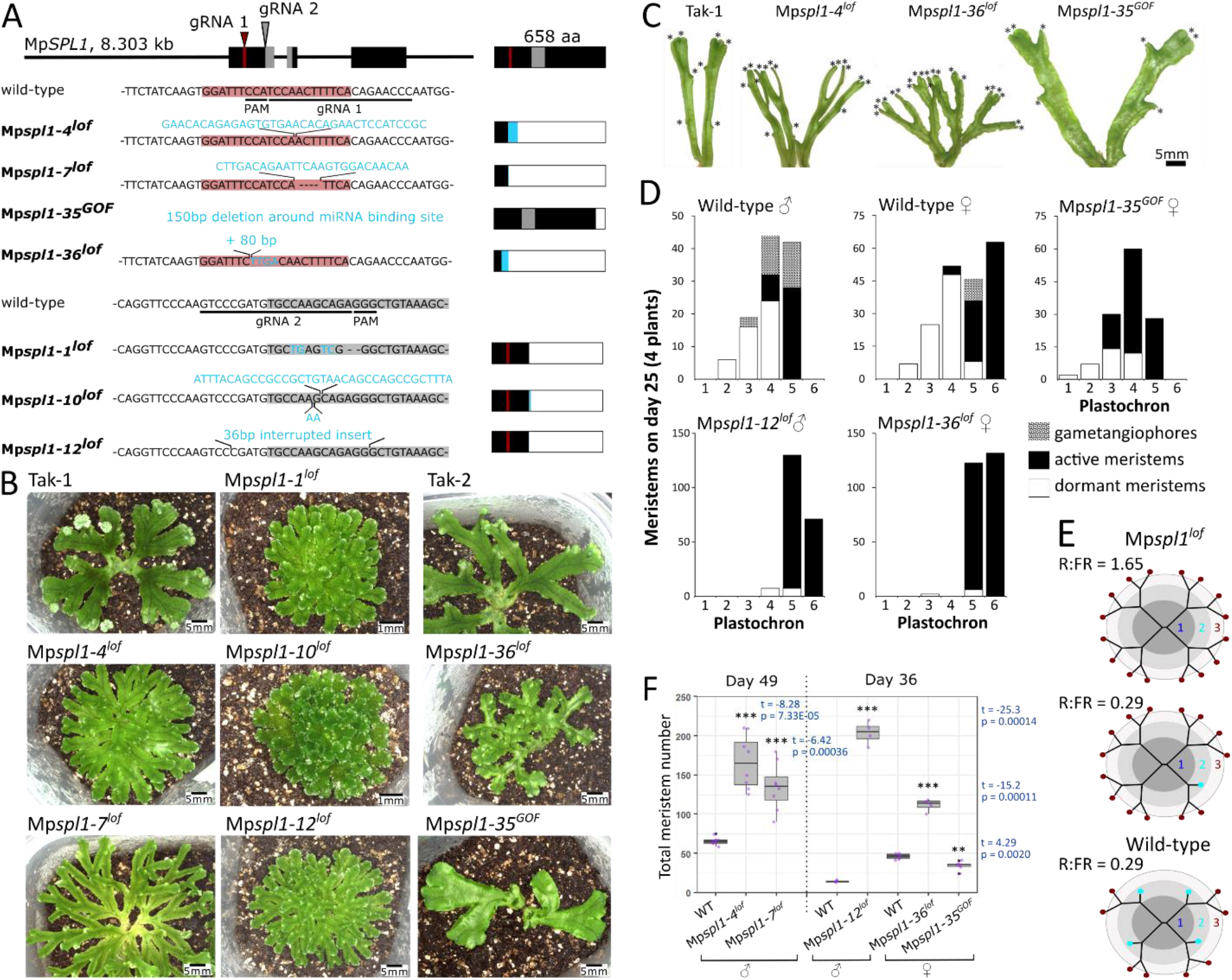
Mp*SPL1* promotes meristem dormancy in early development of the mature thallus. **(A)** Mp*SPL1* gene model with gRNA target sites indicated by arrow heads – gRNA1 in red and gRNA2 in grey. Mpo-MR-13 miRNA binding site is indicated in red, the SBP domain in grey, exons are black boxes. Mutant sequence InDels and SNPs are shown in turquoise colour. Horizontal bars show MpSPL1 protein models with wild-type protein sequence in black, SBP domain in grey, mutant protein sequence in turquoise. White boxes indicate predicted protein truncations due to premature stop codons. Red line indicates integrity of the Mpo-MR-13 miRNA binding site in the MpSPL1 transcript. **(B)** Mp*spl1* mutants grown in low R:FR for 32 days after 14 days of pre-growth in high R:FR light. Tak-1, Mp*spl1-4*^*lof*^, Mp*spl1-7*^*lof*^ (42 days old), Mp*spl1-1*^*lof*^, Mp*spl1-10*^*lof*^ and Mp*spl1-12*^*lof*^ are male lines, Tak-2, Mp*spl1-36*^*lof*^ and Mp*spl1-35*^*GOF*^ (25 days old) are female lines. Scale bars are 5mm except for the ones for Mp*spl1-1*^*lof*^ and Mp*spl1-10*^*lof*^. Male mutants were generated in a Tak-1 wild-type background by thallus transformation, female mutants were generated by transformation of spores obtained by crossing Tak-1 with Tak-2. **(C)** Thallus branches removed from plants grown in low R:FR light as described in (B). Asterisks indicate the positions of meristems (both dormant and active ones). **(D)** Absolute number of gametangiophores (patterned bars), active meristems (black bars) and dormant meristems (white bars) in different plastochrons in Tak-1, female F1 wild-type obtained by crossing Tak-1 with Tak-2, and Mp*spl1*^*lof*^ and Mp*spl1*^*GOF*^ mutants. Plants were grown in low R:FR light for 25 days after 13 days in high R:FR light. For each male line, total meristems of 4 plants were analysed, for each female line, total meristems of 6 plants were analysed. **(E)** Schematics illustrating thallus branching pattern of Mp*spl1*^*lof*^ mutants in high (1.65) and low (0.29) R:FR. Labelling of the schematics is the same as in Figure 1, branching in Mp*spl1*^*lof*^ and wild-type in R:FR = 1.65 is identical (Suppl. Figure 6). **(F)** Average number of meristems per plant in male (Mp*spl1-4*^*lof*^, Mp*spl1-7*^*lof*^, Mp*spl1-12*^*lof*^) and female (Mp*spl1-36*^*lof*^, Mp*spl1-35*^*GOF*^) mutants on day 49 or 36 (data obtained in separate experiments) of growth in low R:FR light after pre-growth in high R:FR for 14 days. Averages between wild-type and mutant were compared using Welch’s t-test, n = 4-8.

**Figure 5:**
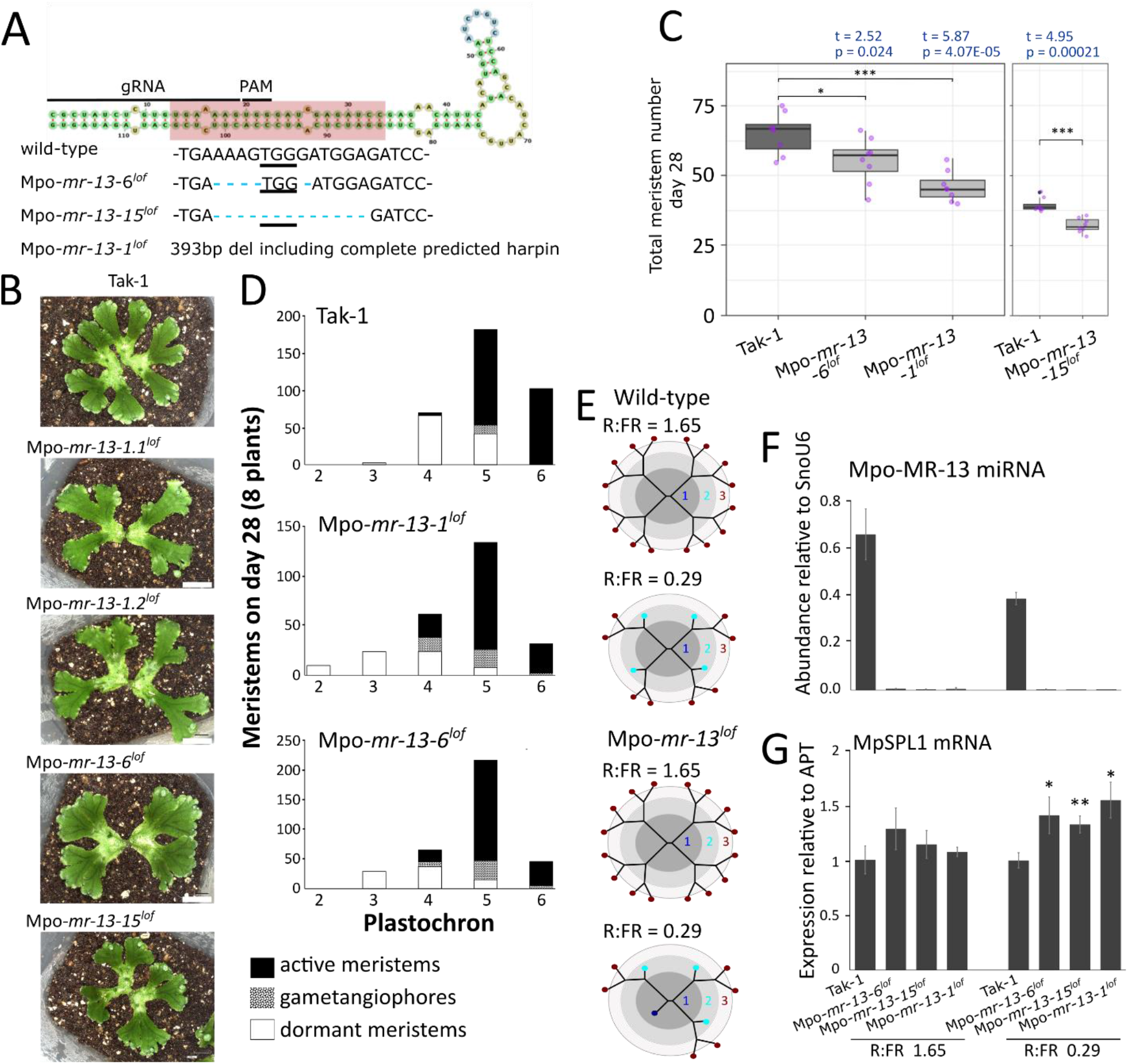
Mpo*-mr-13*^*lof*^ mutants resemble the Mp*spl1*^*GOF*^ mutant. **(A)** Predicted Mpo-MR-13 stem-loop (Tav et al., 2016; Tempel & Tahi, 2012) with 21-nt miRNA sequence highlighted in red and sequence deletions in Mpo*-mr-13*^*lof*^ mutants indicated in light blue. These mutants were generated in a Tak-1 wild-type background. **(B)** Tak-1, Mpo*-mr-13-6*^*lof*^, Mpo*-mr-13-15*^*lof*^ and Mpo*-mr-13-1*^*lof*^ grown in high R:FR (1.65) for 14 days, then in low R:FR light for 21 more days. Mpo*-mr-13-1*.*1*^*lof*^ and Mpo*-mr-13-1*.*2*^*lof*^ are two independent lines of the same allele. Scale bars are 5mm. **(C)** Average total number of meristems per plant grown in low R:FR (0.29) for 28 days. Tak-1 and Mpo*-mr-13-15*^*lof*^ plants were analysed in a separate experiment. In both experiments, gemmae were first grown in high R:FR for 13 days before tranfer to low R:FR light. Averages between Tak-1 and mutants were compared using Student’s t-test with equal variances and n = 8. **(D)** Absolute number of gametangiophores (patterned bars), active meristems (black bars) and dormant meristems (white bars) in different plastochrons in Tak-1, Mpo*-mr-13-1*^*lof*^ and Mpo*-mr-13-6*^*lof*^ mutants grown in high R:FR for 13 days and then in low R:FR for 28 days. Per line, total meristems from 8 plants were analysed. **(E)** Schematics illustrating thallus branching patterns of wild-type and Mpo-*mr-13*^*lof*^ mutants in high (1.65) and (0.29) low R:FR. Labelling of the schematics is the same as in Figure 1. **(F)** Steady state levels of mature Mpo-MR-13 miRNA in Mpo*-mr-13*^*lof*^ mutants grown in high or low R:FR light for 13 days. Mpo-MR-13 miRNA levels were quantified relative to the snoU6 small RNA using qRT-PCR with TaqMan probes. **(G)** Steady state levels of MpSPL1 mRNA relative to MpAPT mRNA in Mpo*-mr-13*^*lof*^ mutants grown in high or low R:FR light for 13 days, measured by qRT-PCR. For each of the two light conditions, steady state levels in mutants were calculated relative to those in Tak-1 wild-type and compared using Student’s t-test with p<0.05*, p<0.01** and p<0.001***, n = 3.

To identify miRNA-targeted Mp*SPL* genes, we first identified and classified Mp*SPL* genes in the *M. polymorpha* genome. *M. polymorpha* encodes a minimal set of four Mp*SPL* genes compared to 15 At*SPL* genes in *A. thaliana* (Tsuzuki et al., 2016). We characterised the evolutionary relationship of Mp*SPL* genes to *SPL* genes from other land plants. Sequence data bases were searched using BLASTp with 15 AtSPL protein sequences as queries. *SPL* genes from the liverwort *M. polymorpha* (4 Mp*SPL* genes), the hornwort *A. agrestis* (4 Aarg*SPL* genes), the moss *P. patens* (12 Pp*SPL* genes) and the angiosperm *Amborella trichopoda* (10 Amtr*SPL* genes) were retrieved. A gene tree was generated from a multiple sequence alignment of conserved sequences in the region of the SBP domain (Figure 3A). There is a single Mp*SPL* gene in each of the three clades defined by monophyletic groups – Mp*SPL1* is in clade III, Mp*SPL4* in clade I and Mp*SPL3* in clade II. The paraphyletic clade IV includes Mp*SPL2*. Next, we identified which of the four MpSPL mRNAs were targeted by a miRNA. There are conserved blocks of sequences in Mp*SPL1* and Mp*SPL2* coding sequences that are complementary to miRNA sequences in databases, suggesting that they are subject to regulation by miRNAs. Mpo-MR-13 is complementary to 21 nucleotides in the predicted MpSPL1 mRNA and miR529c is complementary to a 21-nucleotide sequence in MpSPL2 mRNA (Tsuzuki et al., 2016, 2019a). Furthermore, the MpSPL1 mRNA is cleaved by miRNA Mpo-MR-13 (Tsuzuki et al., 2016). The Mpo-MR-13 miRNA target site is present in *SPL* gene sequences in the Jungermanniopsida and Marchantiopsida liverworts but absent from *SPL* gene sequences in every other lineage of land plants, indicating that the Mpo-MR-13 miRNA is specific to these two classes of liverworts (Figure 3B-C). Mp*SPL2* promotes the far-red light-dependent reproductive transition but not branching under negative control of miR529c, and therefore was not pursued further (Tsuzuki et al., 2019). Based on these data, it was hypothesized that Mp*PIF* controls apical dominance by repressing the expression of Mpo-MR-13.

### Steady state Mpo-MR-13 miRNA and Mp*SPL1* mRNA levels are regulated by Mp*PIF*

To test the hypothesis that Mp*PIF* controls apical dominance by repressing the expression of Mpo*-MR-13* we searched for characteristic PIF-binding sequences in the Mpo-MR-13 promoter. AtPIF proteins bind G-box (5′-CACGTG-3′) and PBE-box (5′-CACATG-3′) motifs in MIR156 promoters and repress transcription in *A. thaliana* (Xie et al., 2017). We found a PBE-box motif in the 5kb-promoter region of Mpo*-MR-13* (Figure 3D) consistent with the hypothesis that transcription of Mpo*-MR-13* is inhibited by MpPIF in low R:FR light, when MpPIF is active. To further test the hypothesis that MpPIF represses expression of Mpo*-MR-13*, we measured Mpo-MR-13 pri-miRNA levels in the Mp*pif-3*^*lof*^ mutant by qRT-PCR in low R:FR. Steady state levels of pri-Mpo-MR-13 were slightly higher in Mp*pif-3*^*lof*^ than in wild-type (Figure 3E), which is consistent with Mp*PIF* inhibiting the expression of Mpo-MR-13 during wild-type development in low R:FR light where MpPIF is active. Furthermore, steady state MpSPL1 mRNA levels were lower in Mp*pif-3*^*lof*^ than in wild-type, consistent with the hypothesis that Mp*PIF* promotes MpSPL1 activity by repressing Mpo-MR-13 expression. Mp*PIF*-mediated repression of Mpo-MR-13 expression and the resulting promotion of Mp*SPL1* expression in low R:FR light suggests that Mp*SPL1* promotes meristem dormancy in plants grown in low R:FR light.

### Mp*SPL1* promotes meristem dormancy

To test the hypothesis that Mp*SPL1* promotes meristem dormancy in low R:FR, we determined the branching phenotype of mutants with defective Mp*SPL1* function. We predicted that dormancy would not develop in Mp*spl1* loss-of-function mutants, and each meristem generated by apical meristem duplication would elongate to form a branch axis. Six Mp*spl1* loss-of-function mutants and one putative gain-of-function mutant were generated using CRISPR/Cas9 with two independent guide RNAs (gRNAs). gRNA 1 targeted the Mpo-MR-13 miRNA binding site upstream of the SBP domain sequence, and gRNA 2 targeted the start of the SBP domain sequence (Figure 4A). The Mp*spl1-4*^*lof*^, Mp*spl1-7*^*lof*^ and Mp*spl1-36*^*lof*^ mutants generated with gRNA 1 had premature stop codons upstream of the SBP domain sequence, generating predicted proteins of 48-92 amino acids instead of 658 amino acids in wild-type (Figure 4A). We predict that loss of the SBP domain abolishes transcription factor function (Birkenbihl et al., 2005). Further loss-of-function alleles, Mp*spl1-1*, Mp*spl1-10* and Mp*spl1-12* were generated using gRNA 2. In all three cases, premature stop codons resulted in loss of the SBP domain without disrupting the Mpo-MR-13 binding site (Figure 4A). There was an in-frame deletion of 150bp that included the Mpo-MR-13 miRNA binding site in a putative hypermorph, Mp*spl1-35*^*GAIN OF FUNCTION (GOF)*^ (Figure 4A). Consequently, the predicted mRNA lacked the target for the Mpo-MR-13 miRNA and the protein was likely a functional transcription factor.

All mature Mp*spl1*^*lof*^ mutant plants were more branched than wild-type when grown in low R:FR light conditions for 32 days (Figure 4B, 4C, 4E). There were no dormant meristems in the mutants while wild-type developed dormancy in meristems formed in plastochrons 2, 3, 4 and 5 after 25 days of growth (Figure 4D). Every new meristem that was formed grew out into a branch axis in Mp*spl1*^*lof*^ mutants. The total meristem number per plant was significantly higher in Mp*spl1*^*lof*^ mutants than in wild-type because no meristems were dormant (and each active meristem gives rise to two new meristems while a dormant meristem does not form any new meristems) (Figure 4C, 4F). These data demonstrate that Mp*SPL1* is necessary for meristem dormancy. The Mp*spl1-35*^*GOF*^ mutant that lacked the Mpo-MR-13 binding site developed dormancy earlier than wild-type. Dormancy developed in meristems formed in plastochron 1 in the Mp*spl1-35*^*GOF*^ mutant, while dormancy was never observed in plastochron 1 in wild-type (Figure 4D). As a result, Mp*spl1-35*^*GOF*^developed fewer dominant branches and total meristems than wild-type (Figure 4E). The absence of dormancy in Mp*spl1* loss-of-function mutants and the development of premature dormancy in the putative Mp*spl1* gain-of-function mutant are consistent with the hypothesis that Mp*SPL1* is necessary for meristem dormancy during the development of the thallus branching network.

### Mpo-MR-13 represses Mp*SPL1*-promoted meristem dormancy

Since Mp*SPL1* promotes meristem dormancy and is repressed by Mpo-MR-13, we hypothesized that Mpo-MR-13 would repress meristem dormancy in low R:FR light. To test this hypothesis, we generated three Mpo*-mr-13* loss-of-function mutants in the Tak-1 wild-type background using CRISPR/Cas9. Mpo*-mr-13-6*^*lof*^ and Mpo*-mr-13-15*^*lof*^ had 5bp and 13bp of the miRNA sequence deleted, respectively (Figure 5A). We predicted that these deletions prevent binding of the mature Mpo-MR-13 miRNA to the MpSPL1 mRNA. The third allele, Mpo*-mr-13-1*^*lof*^, contained a 393bp deletion of the complete predicted hairpin structure. Using TaqMan probes complementary to the wild-type Mpo-MR-13 sequence
(TGAAAAGTGGGATGGAGATCC), Mpo-MR-13 miRNA was amplified from the wild-type background, but not from Mpo*-mr-13-1*^*lof*^, Mpo*-mr-13-6*^*lof*^ or Mpo*-mr-13-15*^*lof*^ (Figure 5F). This indicates that mature Mpo-MR-13 miRNA did not accumulate in any of the three Mpo*-mr-13*^*lof*^ mutants. We hypothesized that Mpo-MR-13-mediated MpSPL1 mRNA slicing would not occur and steady state levels of MpSPL1 mRNA would be higher in Mpo*-mr-13*^*lof*^ mutants than in wild-type plants when grown in low R:FR light. Steady state levels of MpSPL1 mRNA in 14-day-old Mpo*-mr-13*^*lof*^ plants quantified by qRT-PCR were higher in *Mpo-mr-13*^*lof*^ than in wild-type in low but not in high R:FR light (Figure 5G). These data are consistent with the hypothesis that Mpo-MR-13 represses Mp*SPL1* in low R:FR light.

We hypothesized that the Mpo-MR-13 miRNA promotes branching by repressing dormancy because loss-of-function mutants in its target Mp*SPL1* develop more branches than wild-type. Mpo*-mr-13*^*lof*^ mutants grown in low R:FR light for 28 days formed more dormant meristems that were arrested in plastochron 2 and 3 than Tak-1 wild-type (Figure 5D) and therefore fewer total meristems per plant than wild-type (Figure 5C). This suggests that Mpo-MR13 represses meristem dormancy and promotes branching. Furthermore, the phenotype of the Mpo*-mr-13*^*lof*^ mutants resembled the Mp*spl1-35*^*GOF*^ phenotype; both genotypes develop more dormant meristems in early plastochrons and therefore fewer branches than wild-type (Figure 5B, 5E). Consequently, both genotypes form a thallus network that is less branched than wild-type (Figure 5B, 5E). This result is consistent with the hypothesis that Mpo-MR-13 promotes thallus branching and inhibits meristem dormancy in low R:FR light by repressing Mp*SPL1* activity.

### Meristem dormancy during the development of the *M. polymorpha* thallus is regulated by apical dominance

Apical dominance controls the branching architecture of angiosperm shoot systems by imposing dormancy on lateral meristems that develop behind the shoot apex. To test the hypothesis that bud dormancy is controlled by apical dominance in the dichotomously branching *M. polymorpha* thallus, we surgically removed the apices of actively growing axes. (Figure 6B, Suppl. Figure 7). If apical dominance operates, excision of an active meristem should result in the outgrowth of dormant meristems near the site of excision. Three weeks after surgical removal of the apex of a growing axis, the dormant meristem nearest the decapitated apex was activated and developed into a branch axis in all 27 (100%) decapitated axes. In 3 of the 27 decapitated axes, other dormant meristems on the same axis were activated and grew out as branches (Figure 6B). Each new branch axis that developed from the activated meristems underwent two further bifurcation events during the course of the experiment. In control axes that were not surgically decapitated, all dormant meristems remained dormant (Figure 6B). The demonstration that the growing apex of *M. polymorpha* imposes meristem dormancy and suppresses the outgrowth of other branches on the same axis is consistent with the hypothesis that apical dominance modulates branching in the *M. polymorpha* thallus.

**Figure 6:**
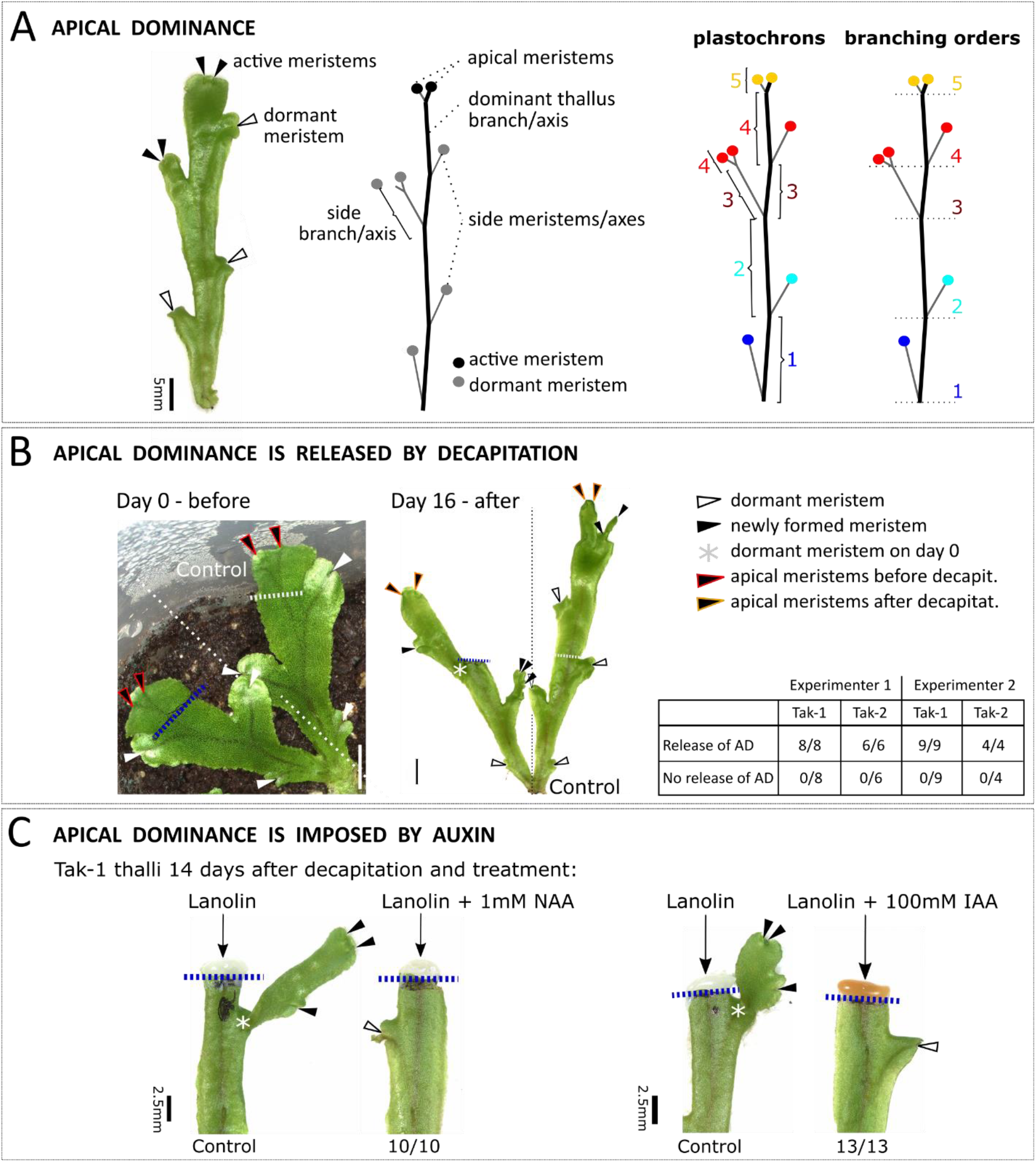
*M. polymorpha* thallus forms apical dominance mediated by Auxin. **(A)** Thallus branches show apical dominance whereby apical meristems are active and side meristems are dormant. The state of each meristem can be described by its plastochron or by its branching order. **(B)** Surgical removal of the apical meristems on the dominant axis causes release of meristem dormancy by day 16 post-decapitation. Side meristems that were dormant on day 0 have grown out and formed new meristems by day 16, including the new apical meristems. Blue broken lines indicate position of the cut, broken grey lines indicate theoretical cut site on the non-decapitated control branch. Two independent experimenters achieved the same result: decapitation releases side branch inhibition in 100% of experiments. **(C)** Tak-1 thallus apices were surgically removed and lanolin containing 1mM NAA or 100mM IAA was applied to the fresh cut site. Controls were treated with pure lanolin. Images show thallus branches 14 days after decapitation and treatment. Application of 1mM NAA or 100mM IAA prevents the outgrowth and division of dormant meristems that occurs in the controls in 10/10 and 13/13 treated thalli. Marks are explained in (B).

Apical dominance is modulated by polar auxin transport from the apical meristem in angiosperms (Leyser, 2005). Auxin produced in the apex moves through the shoot and represses the outgrowth of lateral axes. To test the hypothesis that auxin produced by a dominant meristem modulates apical dominance in *M. polymorpha* by repressing the activity of other meristems and the outgrowth of axes, we surgically removed the apex of the growing branch and applied lanolin paste with or without auxin (Figure 6C). Surgical removal of the apex in controls treated with lanolin paste alone caused the outgrowth of the nearest dormant branch axis. This indicates that lanolin does not have an impact on the release of apical dominance observed when the growing apex is surgically removed. However, the outgrowth of the dormant axis nearest the apex did not occur if lanolin paste containing auxin was applied to the site where the actively growing apex was surgically removed. This indicates that auxin treatment restores the repressive signal that promotes meristem dormancy and inhibits axis outgrowth. Taken together these data are consistent with the hypothesis that apical dominance operates in the dichotomously branching thallus of *M. polymorpha*, and auxin produced by apical meristems imposes meristem dormancy and represses branch axis outgrowth.

Since both Mp*SPL1* and apical dominance promote bud dormancy, we tested if Mp*SPL1* was sufficient to maintain bud dormancy in the absence of repressive auxin signal. If this were the case, we predicted that surgical removal of the apex of the Mp*spl1-35*^*GOF*^ mutant – in which dormancy-promoting Mp*SPL1* activity is higher than in wild-type – would not re-activate dormant meristems. If on the other hand,

MpSPL1 was not sufficient for meristem dormancy, then surgical removal of the apex would result in the activation of dormant meristems. The distal regions of Mp*spl1-35*^*GOF*^ and wild-type thallus apices were surgically removed, and plants were grown for 17 days before the fates of meristems were tested.

Removal of the apices in wild-type plants resulted in the activation of dormant meristems. Removal of the apices in Mp*spl1-35*^*GOF*^ mutant plants also resulted in the activation of dormant meristems (suppl. Figure 8). This indicates that the activation of dormant meristems by surgical removal of the apex is not defective in the Mp*spl1-35*^*GOF*^ mutant. Therefore, we conclude that the control of meristem dormancy by Mp*SPL1* is independent of the control of meristem dormancy by apical dominance.

## DISCUSSION

Two types of branching are found among land plants (Kenrick & Crane, 1997; Hetherington et al., 2020). Dichotomous branching, where the apical meristem duplicates and each daughter meristem grows to form a bifurcation is found among extant bryophyte gametophytes and lycophyte sporophytes (Bierhorst, 1971; Schuster, 1984a; Gola et al., 2014). Lateral branching, where axes branch subapically through the formation of meristems in the axils of determinate structures (leaves) is found in euphyllophyte sporophytes. We show here that light signaling modulates the architecture of the dichotomously branching thallus network of *M. polymorpha* by regulating the activity of meristems produced by bifurcation. The R:FR light ratio modulates phytochrome-regulated expression of the single-copy *M. polymorpha* clade III *SPL* gene, which promotes meristem dormancy by repressing meristem activity. Mp*PIF* controls the expression of the clade III *SPL* gene by modulating the levels of Mpo-MR13, a liverwort-specific miRNA. It has been proposed that PIF-mediated control of MIR 156 in *A. thaliana* modulates the expression of an unidentified member of the clade IV *SPL* genes during the shade avoidance response in *A. thaliana* (Xie et al., 2017). This demonstrates that different miRNAs – Mpo-MR13 in *M. polymorpha* and MIR156 in *A. thaliana –* regulate the expression of different clades of *SPL* genes – clade III in *M. polymorpha* and clade IV in *A. thaliana –* during dichotomous, apical branching *M. polymorpha* and lateral, subapical branching in *A. thaliana*. Taken together these data are consistent with the hypothesis that different mechanisms of miRNA-mediated and SPL-regulated meristem dormancy evolved independently in the liverwort lineage where branching occurs in the gametophyte and in the angiosperm lineage where branching occurs in the sporophyte. If true, the independent recruitment of the different miRNAs and clades *SPL* genes to control dormancy may represent an example of deep homology (Scotland, 2010).

The shade avoidance response is a suite of responses including the repression of lateral meristem development by promoting meristem dormancy in angiosperms. In *A. thaliana* the shade avoidance response is modulated by the red-light receptor phytochrome, which senses the ratio of red to far-red light (R:FR) (Smith & Whitelam, 1997; Lorrain et al., 2008; Casal, 2012). We demonstrate that the F:FR ratio regulates branching architecture by modulating the relative number of active and dormancy meristems in *M. polymorpha* thalli; meristem dormancy is promoted by low R:FR light, characteristic of conditions where light is transmitted through plant tissue or reflected from plant tissue (shade). Furthermore, loss of Mp*PHY* function results in more meristem dormancy than wild-type, while loss of Mp*PIF* function results in the loss of meristem dormancy in plants grown in low R:FR light. Taken together, the modulation of branching by the R:FR ratio, increased meristem dormancy in Mp*phy*, and decreased meristem dormancy in Mp*pif* mutants, are consistent with the hypothesis that apical dominance in *M. polymorpha* is part of a shade avoidance response to limit self-shading.

MpPIF-mediated light signalling regulates meristem dormancy by modulating the expression of the Mp*SPL1* transcription factor (Figure 7). We showed that a single clade III *SPL* gene represses meristem outgrowth and promotes dormancy. Mp*SPL1* is required for meristem dormancy; there is less dormancy in Mp*spl1* mutants than in wild type, and meristem dormancy develops earlier in Mp*spl1* gain-of-fuction mutants than in wild type. The expression of Mp*SPL1* is directly regulated by the Mpo-MR13 miRNA, which targets the MpSPL1 mRNA for slicing. Furthermore, the expression of Mpo*-MR-13* is in turn regulated by Mp*PIF*; Mp*PIF* represses expression of the miRNA in low R:RF light, which increases MpSPL1 mRNA levels. The data are consistent with the hypothesis that high R:FR light destabilizes MpPIF, which increases Mpo-MR-13 miRNA levels that in turn reduce MpSPL1 mRNA levels which stimulates meristem activity and branch outgrowth. By contrast in low R:FR light, MpPIF accumulates and reduces Mpo-MR-13 miRNA levels, which increases MpSPL1 mRNA levels. MpSPL1 activity in turn represses meristem activity and branch outgrowth. Taken together our data demonstrate that a Mp*PIF*–Mpo-MR-13–Mp*SPL1* network controls meristem activity and branch outgrowth in the dichotomous branching system of the *M. polymorpha* thallus (Figure 7).

**Figure 7:**
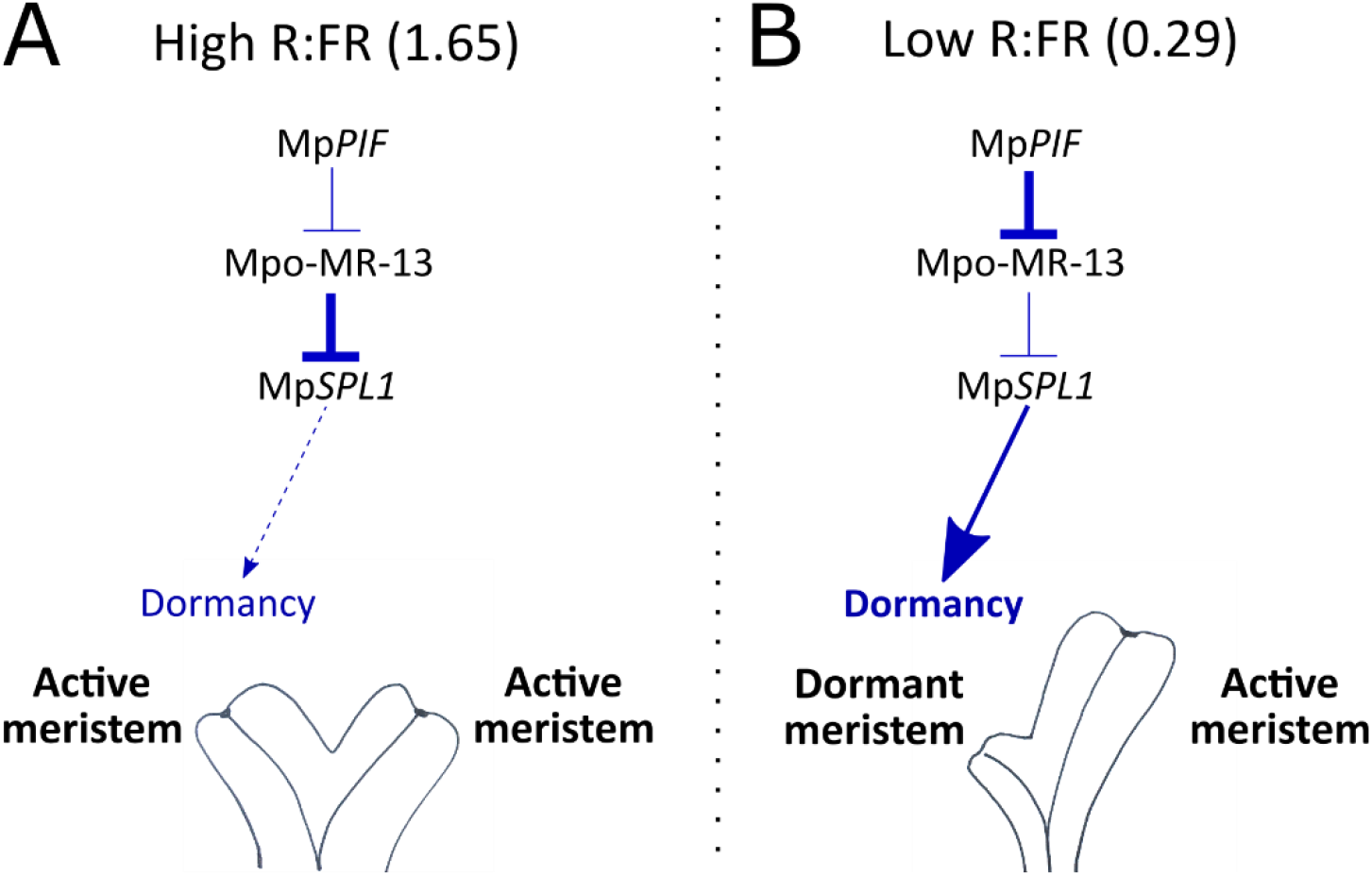
Schematic of the proposed genetic pathway regulating thallus meristem dormancy during apical dominance. Drawn is the shape of a thallus lobe that has developed after apical meristem duplication in high R:FR (left panel) or low R:FR (right panel) light conditions. Arrow indicates a positive effect, the upside down ‘T’ indicates a negative interaction. **(A)** In high R:FR light, MpPIF is inactive (Inoue et al., 2016) and does not repress the expression of Mpo-*MR-13*. High levels of Mpo-MR-13 miRNA are produced, which causes repression of its target Mp*SPL1*. Without MpSPL1 function, no dormancy is induced in daughter meristems after apical meristem duplication. Both active daughter meristems grow out into branches. **(B)** In low R:FR light, MpPIF is active (Inoue et al., 2016) and represses the expression of Mpo-MR-13. Without suppression by the Mpo-MR-13 miRNA, MpSPL1 accumulates and induces dormancy in one of two daughter meristems after apical meristem duplication. Only the active meristem grows out into a branch, while the dormant meristem does not.

There is evidence that the repression of branching by clade III SPL genes may be conserved among the bryophytes. The moss, *Physcomitrium patens*, develops gametophores – shoot like structures – from branches that form on filamentous protonema (Riese et al., 2008). Branches develop from sub-apical cells that divide obliquely to form a bud which forms the gametophore (Coudert et al., 2015). *P. patens* mutants with reduced clade III *SPL* gene function develop more side branches than wild-type, indicating that light-regulated protonemal branching is repressed by clade III Pp*SPL* genes in the moss (Riese et al., 2008). While the development of moss protonema and the liverwort thallus are very different, the role of clade III *SPL* genes in negatively regulating branching is consistent with the hypothesis that the function of clade III *SPL* genes in branching repression is conserved between liverworts and mosses. There is no known miRNA that regulates clade III Pp*SPL* genes in *P. patens*, which in consistent with the hypothesis that the miRNA that targets class III SPL mRNA in liverworts was not present in the common ancestor but evolved in the lineage giving rise to extant liverworts.

## Conclusion

We show that clade III SPL genes promote meristem dormancy in light-regulated development of the dichotomously branching *M. polymorpha* thallus network. The role of clade III *SPL* genes in the repression of meristem activity is conserved among bryophytes while a different clade of SPL genes – clade IV *SPL* genes – may carry out a similar function in *A. thaliana* and other angiosperms. Furthermore, different miRNAs regulate the expression of these different *SPL* genes; Mpo-MR13 regulates *M. polymorpha* clade III *SPL* activity and MIR156 regulates *A. thaliana* clade IV *SPL* activity. Taken together, these data demonstrate that miRNA-mediated *SPL* gene regulation of meristem dormancy evolved independently in the liverwort lineage and in the angiosperm lineage.

## MATERIALS and METHODS

### Plant material and growth conditions

*M. polymorpha* accessions Takaragaike-1 (Tak-1, female) and Takaragaike-2 (Tak-2, female) (Ishizaki et al., 2016) were used as wild type plants for transformations and crossings. Plants were cultivated on ½ Gamborg medium supplemented with 1% agar, in 8 h of darkness alternating with 16 h of white light with a red:far-red light ratio (R:FR) of 1.65. Sexual reproduction was induced by far-red-enriched irradiation (R:FR = 0.29) as described in Chiyoda et al 2008, at 23 °C under a light:dark cycle of 16:8 hours.

### Phylogenetic analyses

Gene trees were built from multiple sequence alignments of amino acid sequences generated on the MAFFT online website. Amino acid and nucleotide sequences from *SPL* genes were identified by BLASTp search with AtSPL protein sequences from the TAIR website as queries. Orthologous protein and nucleotide sequences in *M. polymorpha, P. patens, A. trichopoda, P. trichocarpa* and *S. purpurea* were identified by BLASTp search on the phytozome JGI website (Suppl. Table 1). SPL orthologs in other liverwort species were identified by BLASTp of MpSPL proteins against the 1000 plants (1KP) genome sequence database. SPL protein and nucleotide sequences in the hornwort *A. agrestis* were identified in a local BLASTp and tBLASTn search using the annotated *A. agrestis* proteome or genome from Zurich University (https://www.hornworts.uzh.ch/en/download.html). Multiple sequence alignments were manually trimmed to remove large gaps. Gene trees were built in MEGA-X using the Maximum likelihood statistical method with 1000 (class III *SPL* gene tree) or 500 (*SPL* gene tree) bootstrap resampling steps.

### Phenotyping in low R:FR light (R:FR = 0.29)

4-8 gemmae per line were grown on solid ½ Gamborg medium containing 1% sugar and 1% Agar in constant white light (R:FR = 1.65) for 11-14 days. Plants were then transferred to soil (Neuhaus® Huminsubstrat from Klasmann-Deilmann GmbH:vermiculite 2:1) inside closed microbox containers (produced by SacO_2_, OV80+OVD80, #40 green filter, Non gamma irradiated, autoclavable (NG/NP)) with an air filter and grown in far-red-enriched conditions (R:FR = 0.29). To determine reproductive timing, the number of days of growth in far-red-enriched light was counted until the development of the first gametangiophore bud per plant.

Total meristem number was determined by counting all thallus meristems including dormant and active ones and such that had terminated in a gametangiophore. We illustrated the data using boxplots. Differences between means were determined by student’s t-test when equal sample sizes and Welch’s t-test when unequal sample sizes, assuming unequal variances and p<0.05*, p<0.01**, p<0.001***.

The fate of a thallus meristem was designated ‘dormant’ when after duplication, this daughter meristem did not elongate and duplicate itself to give rise to another branch while the other daughter meristem did. The fate of this second daughter meristem was then ‘active’. Meristem fate was designated ‘gametangiophore’ as soon as a small gametangiophore bud was visible at the site of a thallus meristem.

### CRISPR/Cas9 mutagenesis

Guide RNAs (gRNAs) for CRISPR/Cas9 mutagenesis of Mp*PHY*, Mp*PIF*, Mp*SPL1* and Mpo-MR-13 were designed using the CRISPR-P online tool (Table 1). To make two Mp*SPL1*-targeting and one Mpo-MR-13-targeting single gRNA constructs, we used gateway cloning to recombine an entry vector carrying one gRNA driven by a pMpU6 promoter (vector pHB453, a MpEN03 vector modified by H. Breuninger), with a destination vector encoding the Cas9 enzyme driven by a CamV35S promoter (pMpGEO010) (Sugano et al 2014). The double gRNA construct for mutagenesis of Mpo-MR-13 was cloned using the Golden Gate vectors OP-74, OP-75, OP-005, OP-63 and OP-73 from the Open Plant toolkit and following the corresponding CRISPR cloning protocol (Sauret-Güeto et al., 2020). To make the CRISPR constructs to target Mp*PHY* and Mp*PIF*, we performed loop assembly of the single gRNA oligo duplex and the OP-76 plasmid following the corresponding Golden Gate cloning protocol from (Sauret-Güeto et al., 2020).

**Table 1:**
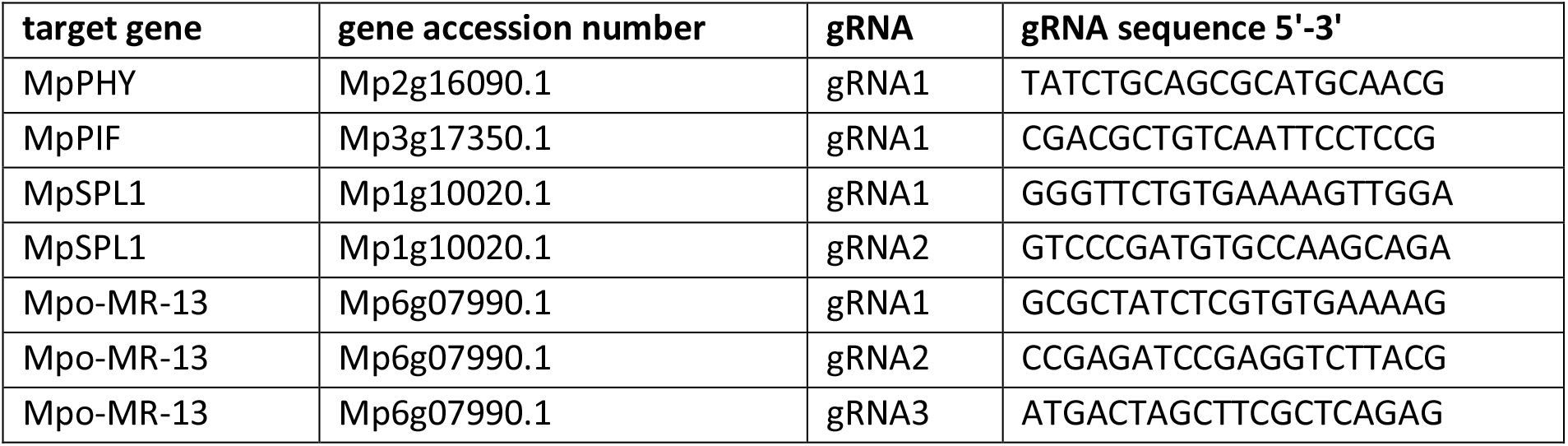
guide RNAs for CRISPR mutagenesis.

We identified primary CRISPR mutants by PCR genotyping, amplifying 50-1000 bp around the gRNA target sites within Mp*PHY*, Mp*PIF*, Mp*SPL1* and Mpo*-MR-13* (Table 2). Gemmae from T1 plants were genotyped for mutations and propagated as isogenic lines.

**Table 2:**
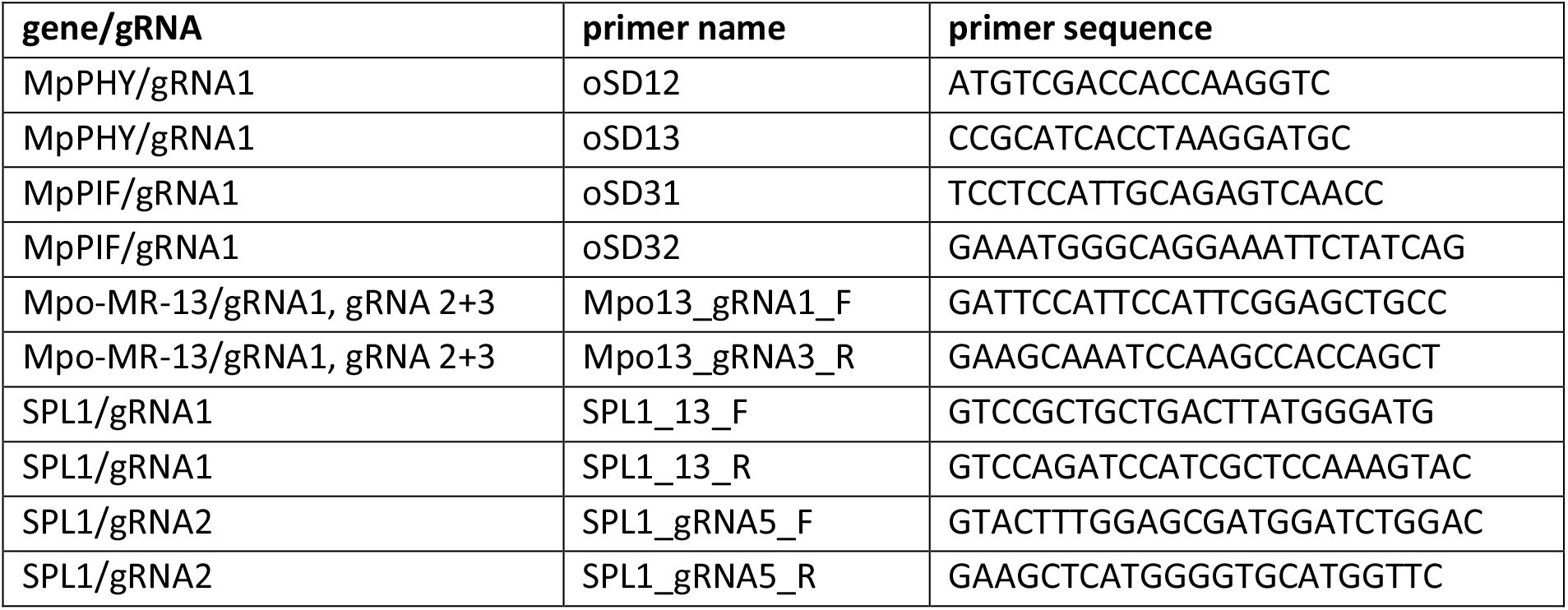
Primers used for PCR genotyping of CRISPR transformants.

### Genotyping of CRISPR mutants

To isolate genomic DNA for PCR genotyping, ~3×3 mm thallus tissue was ground together with 100µl Extraction buffer (100 mM Tris HCl [pH 9.5], 1 M KCl, 10 mM EDTA (Tsuboyama & Kodama, 2014)). The tissue suspensions were incubated at 65 °C for 10 min and diluted 1:5 with sterile Mono Q water. 1µl gDNA template was used in a PCR reaction together with 2x concentrated in-house master mix (developed by Molecular Biology Service) containing hot start polymerase, and 0.5 µM forward and reverse primer, each. PCR was performed with 40 cycles of 95°C for 30 sec, 58°C for 30 sec and 72°C for 1 min. Whole PCR product (20 µl) was treated with 2 µl ExoSap mix containing 10 mM Tris-HCl, 100 mM NaCl, 5 mM ß-Mercaptoethanol, 0.5 mM EDTA, 50% Glycerol, 1 mM MgCl2, adjusted to pH 7.5 at 25°C, and 0.8 U Exonuclease I (NEB) and 0.4 U Shrimp Alkaline Phosphatase (NEB) at 37°C for 30 min. The reaction was stopped at 80°C for 10 min. 1 µl of purified PCR product was sequenced by Sanger sequencing to identify mutations in gRNA target sites.

### Spore and thallus transformation

Male Mp*spl1*^*lof*^ and Mpo*-mr-13*^*lof*^ mutants were generated by Agrobacterium-mediated transformation of regenerated Tak-1 thallus pieces with CRISPR constructs as described in (Kubota et al., 2013). Thallus transformation of Tak-2 failed repeatedly, therefore we transformed F1 spores from a Tak-1 x Tak-2 crossing to generate female Mp*spl1*^*lof*^ and Mp*spl1*^*GOF*^ mutants following the protocol provided in (Chiyoda et al., 2008). The male Mp*phy*^*lof*^ and Mp*pif*^*lof*^ mutants were also generated by spore transformation. Transformed sporelings and thallus pieces were selected on 10 ng µl^-1^ hygromycin.

### Extraction of total RNA and cDNA synthesis

Total RNA was extracted using the RNeasy Mini kit (Quiagen) and following the handbook with one modification – to make RLT buffer, we added 30μl β-Mercaptoethanol per 1ml. We performed on-column DNase I digestion using the RNase-Free DNase set (Quiagen Cat. No. / ID: 79254).

Reverse transcription was performed using the LunaScript RT SuperMix (NEB E3010) and 1μg DNase-treated total RNA with incubation at 25ºC for 2 min, 55ºC for 10 min and 95ºC for 1 min.

### Quantitative real time PCR (qRT-PCR)

To quantify mRNA levels, we performed qRT-PCR using Luna® Universal qPCR Master Mix (M3003). We prepared reactions with volumes of 10 μl by adding 200 ng cDNA and 0.3 µM of each forward and reverse primer (Table 3). We generated standard curves with four dilutions (1:10, 1:100. 1:1000, .1:10.000) for each primer pair. Ser sample, three technical replicates were included. qRT-PCR was performed inside a QuantStudio 7 cycler (Applied Biosystems) with initial denaturation at 95°C for 1 min followed by 40 cycles of 95°C for 15 sec and 60°C for 30 sec.

**Table 3:**
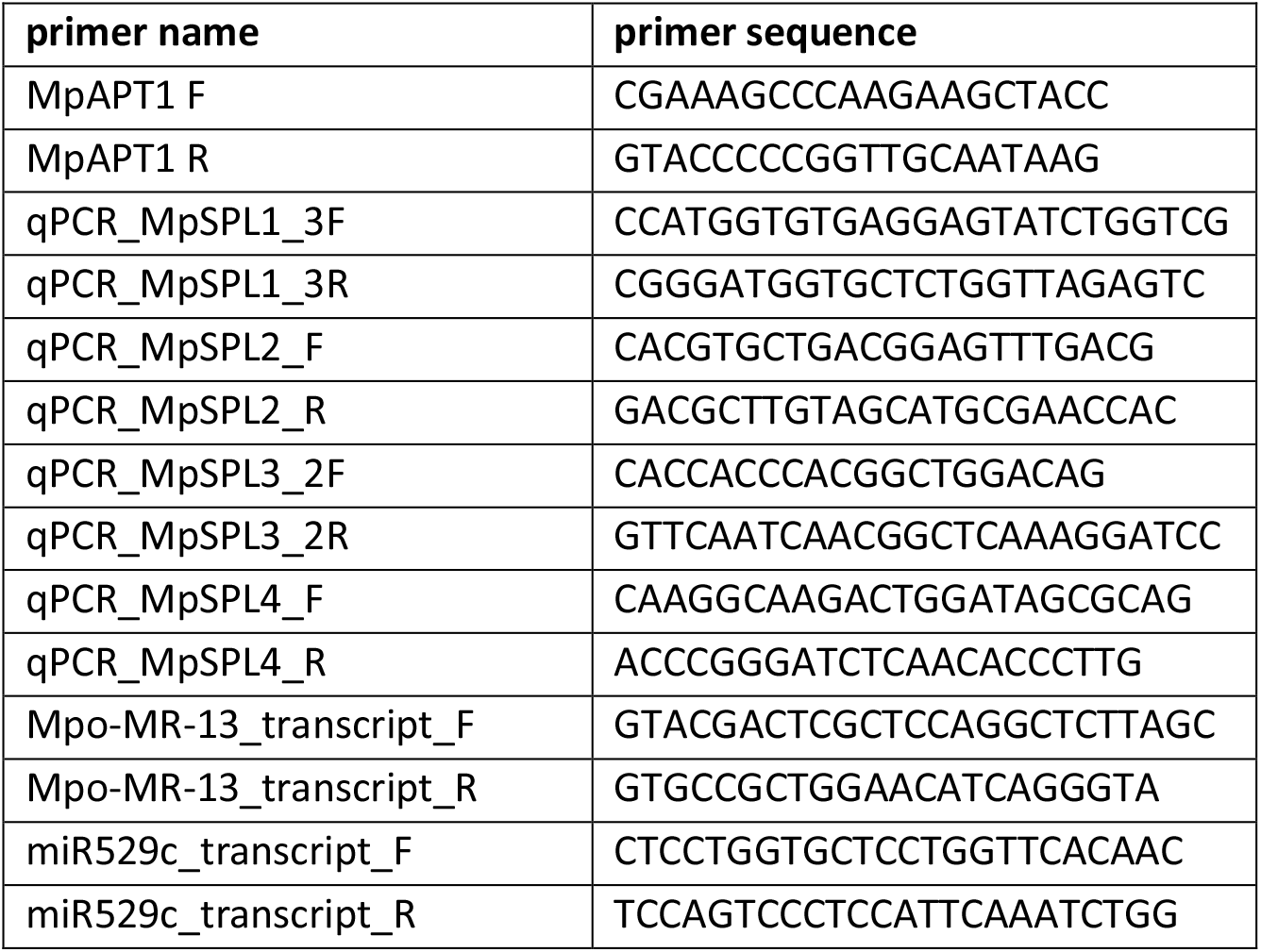
qRT-PCR primers.

The obtained Ct values were evaluated by the Pfaffl method to obtain relative gene expression (Pfaffl, 2007). First, primer efficiencies (E) were determined from the standard curve using the formula 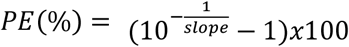 and converted using the formula 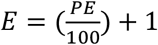 Then, average ΔCt values were calculated for the genes of interest (GOI) and the housekeeping gene APT1 (HKG) by determining the ‘control average’ value for each gene and subtracting each Ct average from it. Finally, the Pfaffl equation was applied to calculate gene expression ratios for each gene of interest: 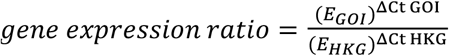 (Pfaffl, 2001).

### TaqMan miRNA quantification assay

Custom TaqMan® probes (including RT primers and qRT-PCR primers, Thermo Scientific) were designed using the Small RNA Assay Design Tool with the following input sequences: UGAAAAGUGGGAUGGAGAUCC for Mpo-MR-13, CCAGAAGAGAGAGAGCACAGC for MpmiR529c and GAGAAGAUUAGCAUGGCCCCU for the MpsnoU6 control. Reverse transcription was performed using the TaqMan™ MicroRNA Reverse Transcription Kit (Thermo Scientific) with 10ng of total RNA template, following the instructions of the TaqMan™ Small RNA Essays user guide. The RT product was diluted 2X with nuclease-free water, and 2μl (1 ng cDNA) used as a template for qRT-PCR. The qRT-PCR reaction contained 20X TaqMan™ probe and 2X TaqMan™ Fast Advanced Master Mix (Thermo Scientific) in a total of 10 μl. qRT-PCR was performed inside a QuantStudio 7 cycler (Applied Biosystems) with enzyme activation at 95°C for 20 sec followed by 40 cycles of 95°C for 1 sec and 60°C for 20 sec.

### Microscopy

Images of whole Marchantia plants, thallus branches and gametangiophores were taken with the Keyance digital bright field microscope VHX-7000 using the 20x-200x objective lens.

**Gene accession numbers** (Supplementary Table 1).

## Notes

### Competing Interest Statement

The authors have declared no competing interest.

