## Supplemental Table and Figures for "Meristem dormancy in a dichotomous branching system is regulated by a liverwort-specific miRNA and a clade III *SPL* gene in *Marchantia polymorpha*"

**SUPPLEMENTARY MATERIAL**

**Supplementary Table 1:** BLAST results and gene accessions used to build *SPL* gene trees and nucleotide sequence alignment.

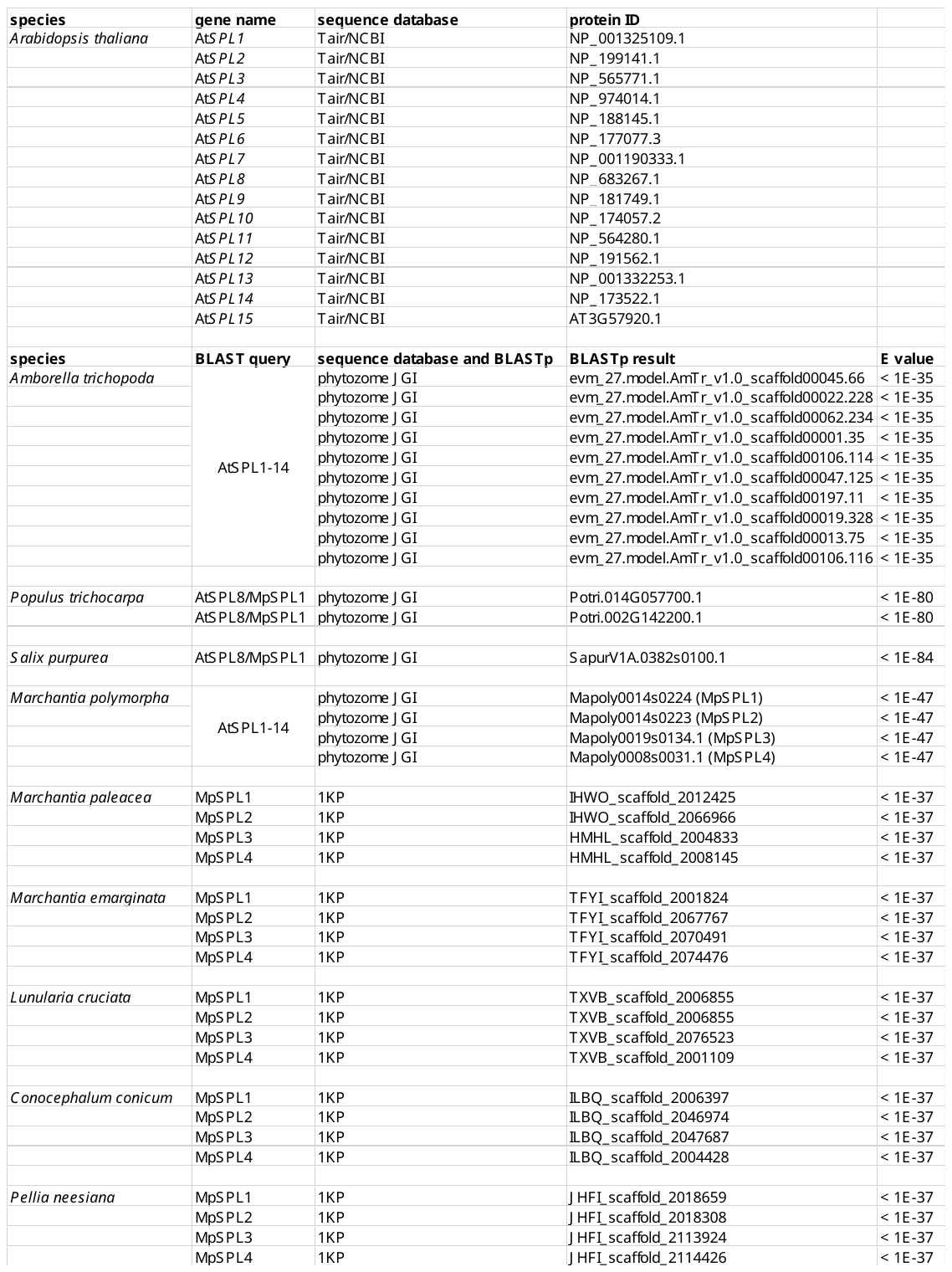

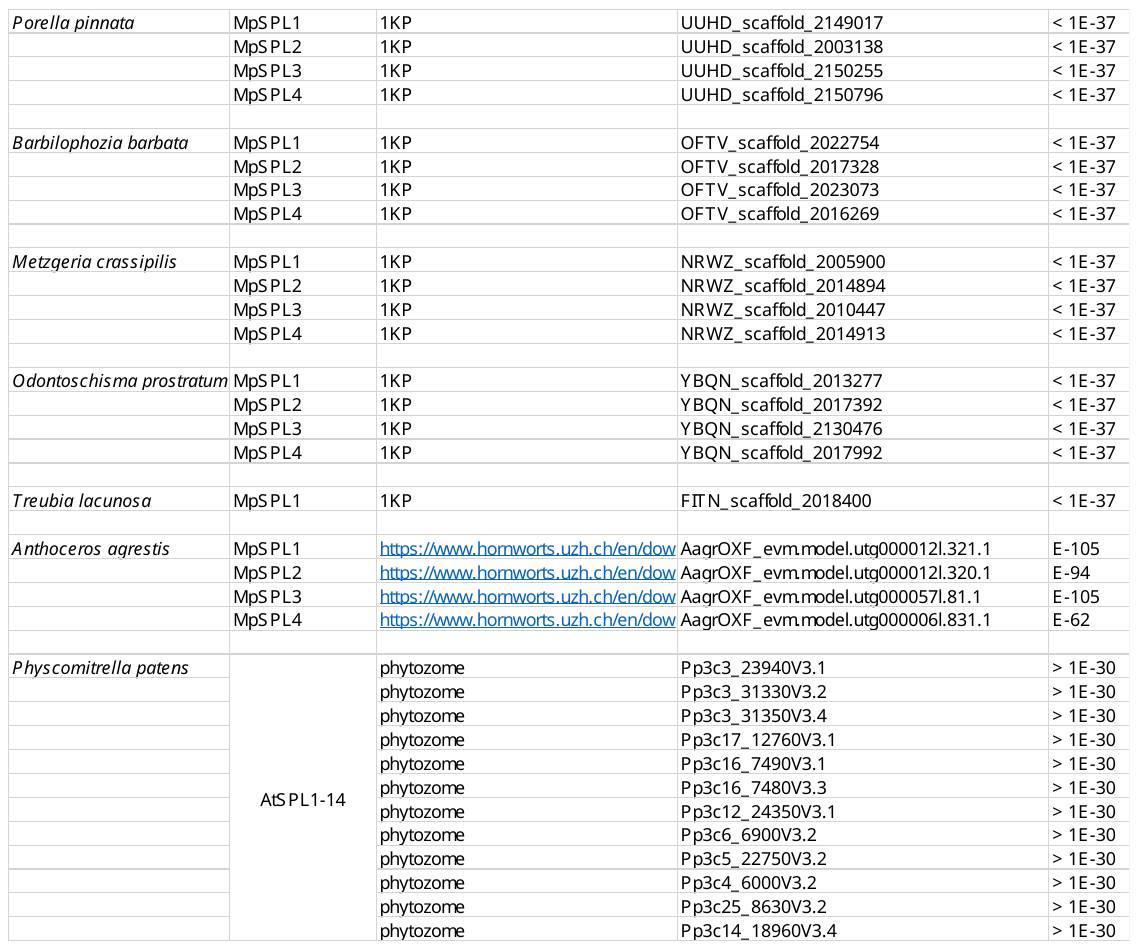

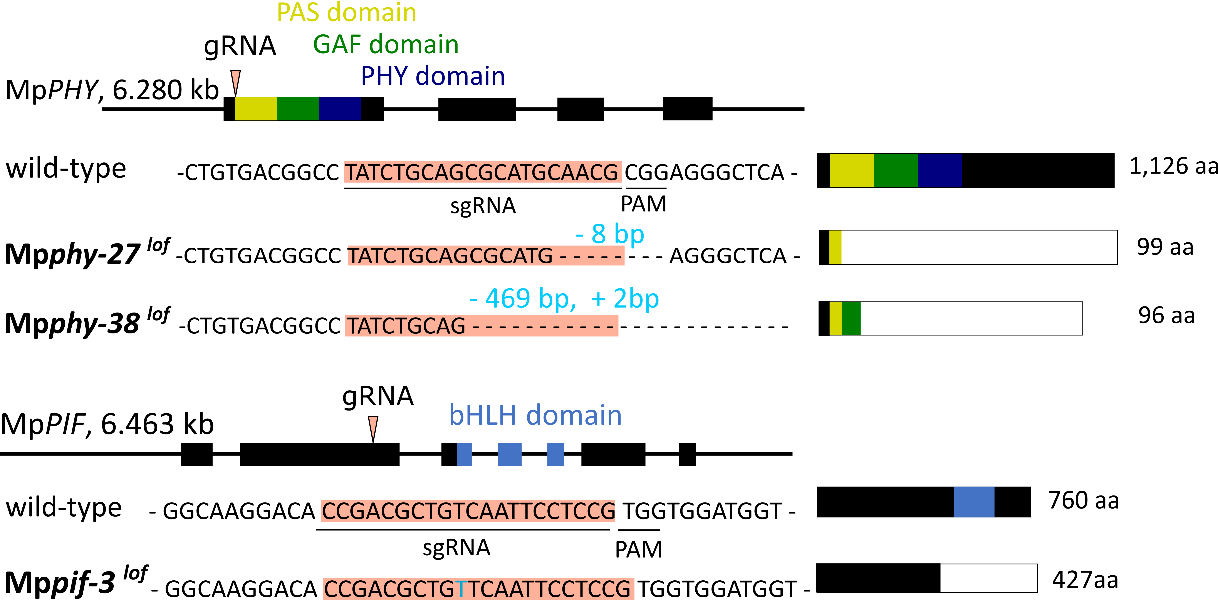

**Supplementary Figure 1: Generation of Mp*phy^lof^* and Mp*pif^lof^* mutants.** Mp*PHY* and Mp*PIF* gene models with gRNA target sites indicated by red arrow heads. Functional domains are indicated in yellow (PAS domain), green (GAF domain) and dark blue (PHY domain) in Mp*PHY*, and in blue (bHlH domain) in Mp*PIF*. Below, nucleotide sequences around the gRNA binding sites (highlighted in red) in wild-type and mutant alleles are shown, with mutant sequence InDels in turquoise colour. Horizontal bars show MpPHY and MpPIF protein models with wild-type protein sequence in black and functional protein domains in the colours described above. White boxes indicate predicted protein truncations due to premature stop codons

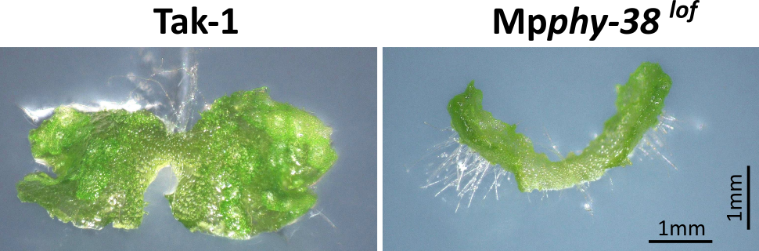

**Supplementary Figure 2: Mp*phy^lof^* mutants constitutively undergo orthotropic growth.** Tak-1 and Mp*phy-38^lof^* gemmae grown on ½ Gamborg medium with 1% sucrose in high R:FR (1.65) for 7 days. Gemmae were imaged at a 45 degrees angle. Mp*phy-38^lof^* thallus bends upwards as visible by the rhizoids that appear white and span between thallus and substrate.

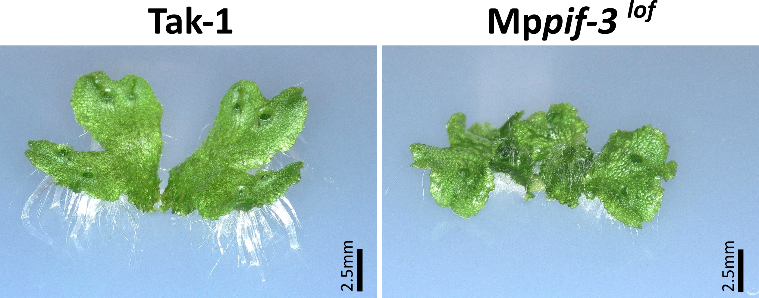

**Supplementary Figure 3:** **Mp*pif^lof^* mutant does not grow orthotopically in response to low R:FR light.** Tak-1 and Mp*pif-3^lof^* gemmae grown on ½ Gamborg medium with 1% sucrose in high R:FR (1.65) for 10 days and then in low R:FR for 6 days. Gammae were imaged at a 55 degrees angle. Tak-1 wild-type thallus bends upwards as visible by the rhizoids that appear white and span between thallus and substrate.

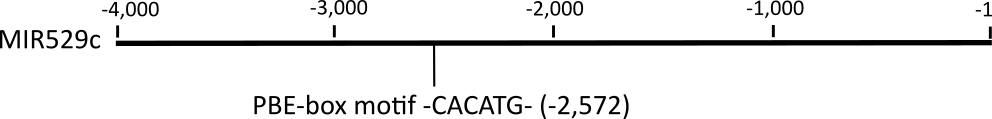

**Supplementary Figure 4: MIR529c contains a putative PIF-binding motif upstream of the miRNA sequence.** PBE-box motif 2,572 bp upstream of the miR529c miRNA sequence.

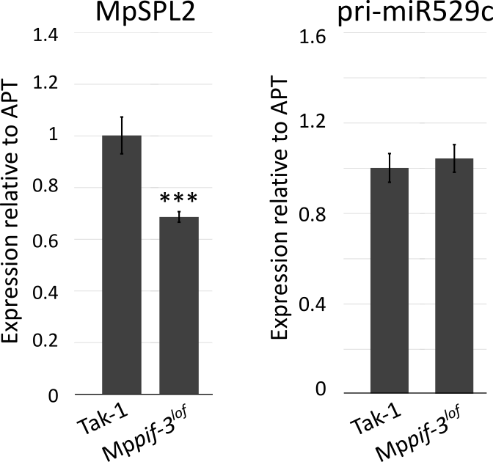

**Supplementary Figure 5: Through repression of miR529c, Mp*PIF* promotes the expression of Mp*SPL2* in low R:FR light.** Steady state levels of MpSPL2 mRNA and pri-miR529c in Tak-1 and Mp*pif-3^lof^* in whole gemmae grown in low R:FR light for 14 days. The insignificant increase in pri-miR529c in Mp*pif-3^lof^* relative to Tak-1 is likely due to the developmental stage of the gemmae, which had not initiated gametangiophores by day 14. A slight increase in pri-miR529c translates into a significant decrease in MpSPL2 mRNA in Mp*pif-3^lof^*.

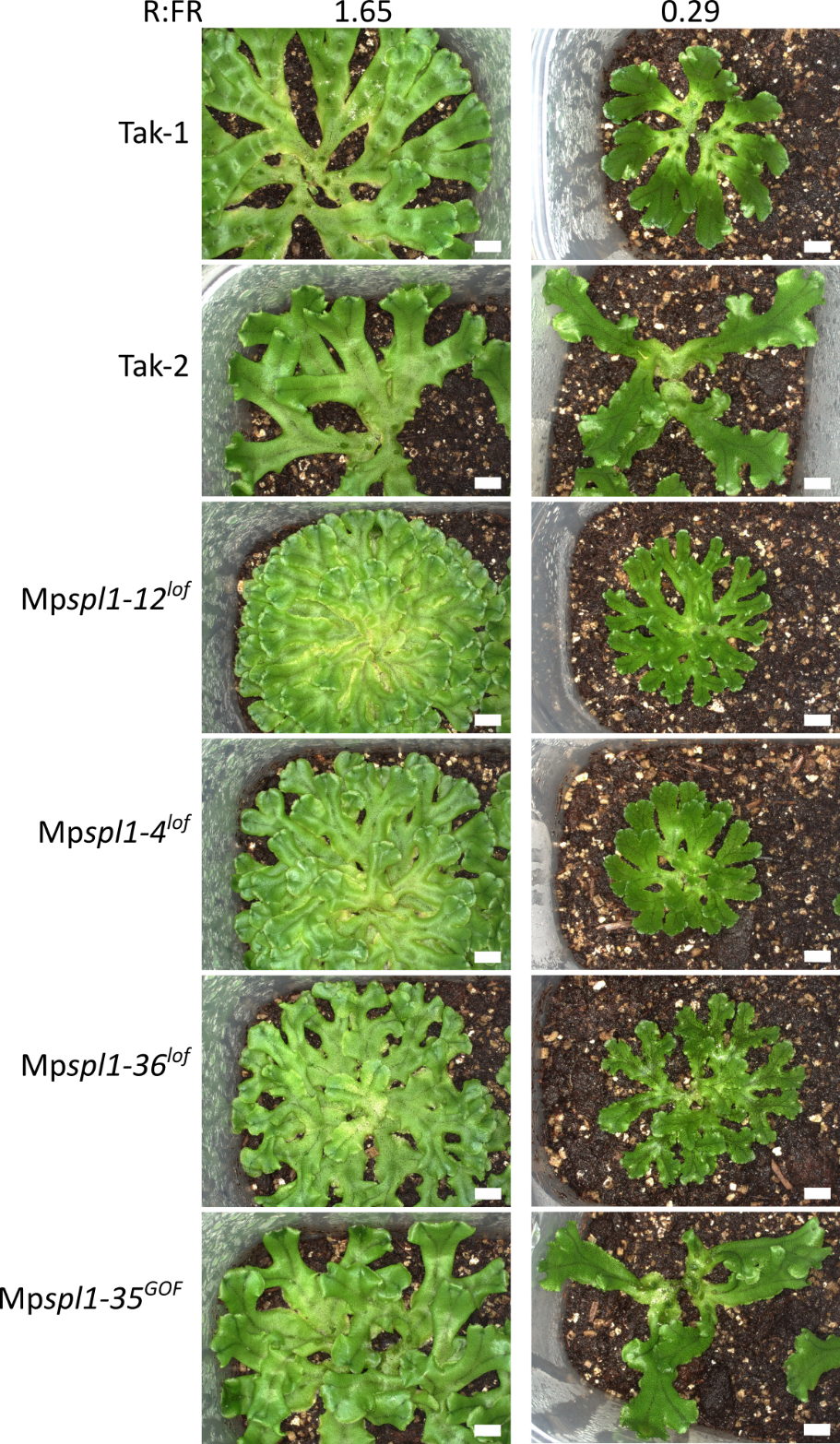

**Supplementary Figure 6: Thallus phenotypes of Mp*spl1^lof^* and Mp*spl1^GOF^* mutants grown in high and low R:FR light.** Plants were grown from gemmae in high R:FR light conditions on ½ Gamborg medium supplemented with 1% Agar for 14 days and then transferred to soil in micropots and grown for another 15 days in either high R:FR (1.65, left panel) or low R:FR (0.29, right panel). Scale bars are 5mm.

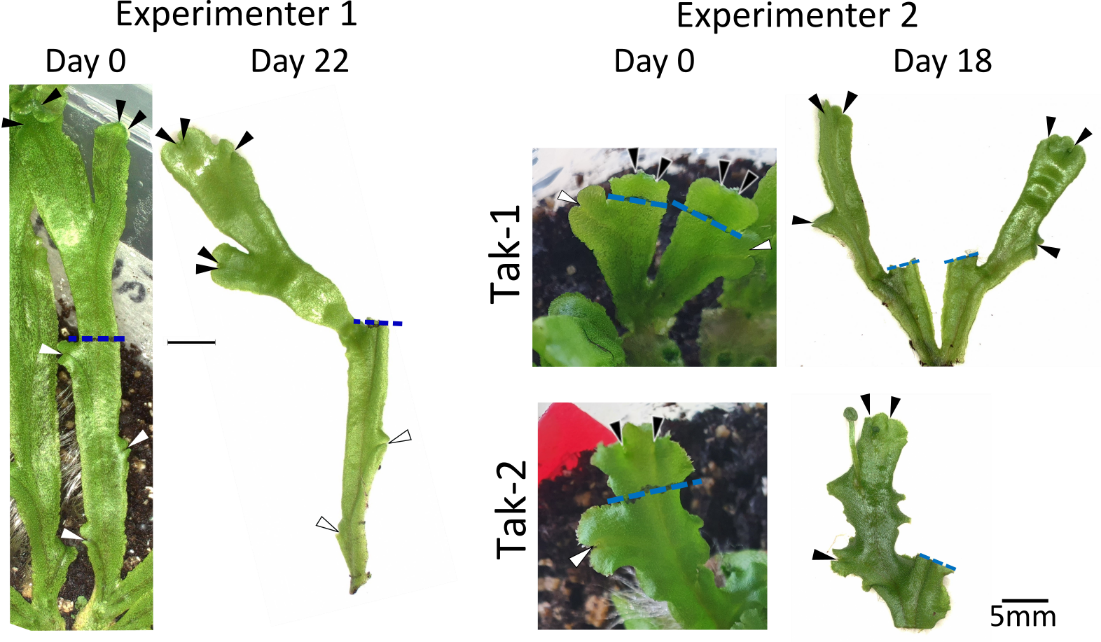

**Supplementary Figure 7: Decapitation of the dominant thallus apex releases apical dominance.** More examples of a decapitation experiments in Tak-1 and Tak-2 wild-type – Decapitation of main branch on day 0 and release of apical dominance by day 22 as performed by experimenter 1, and decapitated thalli on day 0 and day 18 as performed by experimenter 2. Blue broken line indicates position of the cut, white and black arrow heads point at dormant and newly formed meristems, respectively.

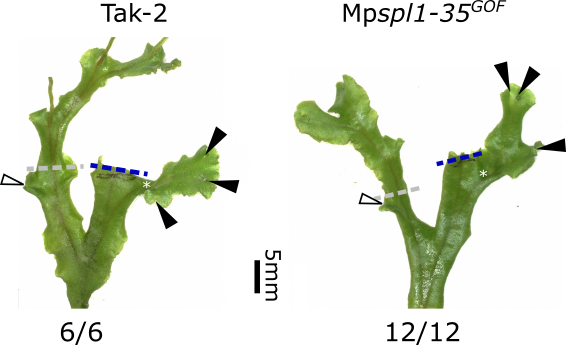

**Supplementary Figure 8: Mp*SPL1* is not sufficient to maintain meristem dormancy upon decapitation of the thallus apex.** Surgical removal of the apices causes release of meristem dormancy in Tak-2 wild-type and the Mp*spl1-35^GOF^* mutant. Side meristems that were dormant on day 0 (indicated by white asterisks on the decapitated branches and by white arrow heads on the control branches) have grown out and formed new meristems (indicated by black arrow heads) by day 17, the time point that is shown here. Blue broken lines indicate positions of the cut, broken grey lines indicate theoretical cut sites on the non-decapitated control branches. In both genotypes, 100% of experiments resulted in dormancy release.
